# Cleavage of α-1,4-Glycosidic Linkages by the Glycosylphosphatidylinositol-Anchored α-Amylase AgtA Decreases the Molecular Weight of Cell Wall α-1,3-Glucan in *Aspergillus oryzae*

**DOI:** 10.1101/2022.10.12.511150

**Authors:** Ami Koizumi, Ken Miyazawa, Makoto Ogata, Yuzuru Takahashi, Shigekazu Yano, Akira Yoshimi, Motoaki Sano, Masafumi Hidaka, Takanori Nihira, Hiroyuki Nakai, Satoshi Kimura, Tadahisa Iwata, Keietsu Abe

## Abstract

*Aspergillus* fungi contain α-1,3-glucan with a low proportion of α-1,4-glucan as a major cell wall polysaccharide. Glycosylphosphatidylinositol (GPI)-anchored α-amylases are conserved in *Aspergillus* fungi. The GPI-anchored α-amylase AmyD in *Aspergillus nidulans* has been reported to directly suppress the biosynthesis of cell wall α-1,3-glucan but not to degrade it *in vivo*. However, the detailed mechanism of cell wall α-1,3-glucan biosynthesis regulation by AmyD remains unclear. Here we focused on AoAgtA, which is encoded by the *Aspergillus oryzae agtA* gene, an ortholog of the *A. nidulans amyD* gene. Similar to findings in *A. nidulans, agtA* overexpression in *A. oryzae* grown in submerged culture decreased the amount of cell wall α-1,3-glucan and led to the formation of smaller hyphal pellets in comparison with the wild-type strain. We analyzed the enzymatic properties of recombinant (r)AoAgtA produced in *Pichia pastoris* and found that it degraded soluble starch, but not linear bacterial α-1,3-glucan. Furthermore, rAoAgtA cleaved 3-α-maltotetraosylglucose with a structure similar to the predicted boundary structure between the α-1,3-glucan main chain and a short spacer composed of α-1,4-linked glucose residues in cell wall α-1,3-glucan. Interestingly, rAoAgtA randomly cleaved only the α-1,4-glycosidic bonds of 3-α-maltotetraosylglucose, indicating that AoAgtA may cleave the spacer in cell wall α-1,3-glucan. Consistent with this hypothesis, heterologous overexpression of *agtA* in *A. nidulans* decreased the molecular weight (MW) of cell wall α-1,3-glucan. These *in vitro* and *in vivo* properties of AoAgtA suggest that GPI-anchored α-amylases can degrade the spacer α-1,4-glycosidic linkages in cell wall α-1,3-glucan before its insolubilization, and this spacer cleavage decreases the MW of cell wall α-1,3-glucan *in vivo*.

## INTRODUCTION

The cell wall of fungi is composed mainly of polysaccharides; it protects cells from stresses and maintains cell morphology (Latgé, 2010; Yoshimi et al., 2016). *Aspergillus* cell wall is composed of α-1,3-glucan (with a low proportion of α-1,4-glucan), β-1,3-glucan (with β-1,6-glucan branches), galactomannan and chitin. (Latgé, 2010; Yoshimi et al., 2016). In pathogenic fungi, α-1,3-glucan conceals cell wall β-1,3-glucan and chitin and consequently prevents recognition by the host immune system (Rappleye et al., 2004, 2007; Fujikawa et al., 2009, 2012; Beauvais et al., 2013). α-1,3-Glucan is also an adhesion factor in hyphal aggregation (Yoshimi et al., 2013; Miyazawa et al., 2016, 2018; Zhang et al., 2017).

Two α-1,3-glucan synthase genes (*agsA, agsB*) are known in the model fungus *Aspergillus nidulans*, with *agsB* functioning mainly during vegetative hyphal growth (Yoshimi et al., 2013). In *A. nidulans*, the *amyD* and *amyG* genes are located near the *agsB* locus and are predicted to encode glycosylphosphatidylinositol (GPI)-anchored α-amylase and intracellular α-amylase, respectively (He et al., 2014). The cluster of these three genes (*agsB*–*amyD*–*amyG*) is conserved among *Aspergillus* fungi except for *Aspergillus fumigatus* (He et al., 2014).

The mechanism of cell wall α-1,3-glucan biosynthesis was first predicted in the fission yeast *Schizosaccharomyces pombe* (Grün et al., 2005). In the cytoplasm, an α-1,3-glucan synthase Ags1 transfers glucose units from UDP-glucose to the non-reducing end of the primer maltooligosaccharide and thus forms an α-1,3-glucan unit, which is then exported by the same enzyme to the extracellular space (Grün et al., 2005). The extracellular domain of Ags1 is thought to connect two exported units by transglycosylation (Grün et al., 2005). In *S. pombe*, a spacer oligosaccharide (≈12 residues), which is derived from the primer oligosaccharide, is present between two α-1,3-glucan chains (≈120 residues each) (Grün et al., 2005). We previously estimated the mechanism of cell wall α-1,3-glucan biosynthesis in *A. nidulans* with reference to the speculative mechanism in *S. pombe* (Yoshimi et al., 2017; Miyazawa et al., 2018, 2020; Figure 1). Cell wall α-1,3-glucan is mainly synthesized by AgsB, which is composed of the extracellular, intracellular, and multitransmembrane domains (Yoshimi et al., 2017; Miyazawa et al., 2020). The primer maltooligosaccharides, which are likely required for α-1,3-glucan biosynthesis, are predicted to be produced by AmyG (He et al., 2014; Miyazawa et al., 2018, 2020). Kazim et al. (2021) reported that, under submerged culture conditions, *amyG* disruption in *A. nidulans* causes the formation of dispersed hyphae and decreases pellet size and α-1,3-glucan content in the cell wall in comparison with the parental strain; these changes can be reverted by adding maltose or maltotriose to the culture media, suggesting that AmyG provides maltooligosaccharides for α-1,3-glucan biosynthesis.

**Figure 1.**
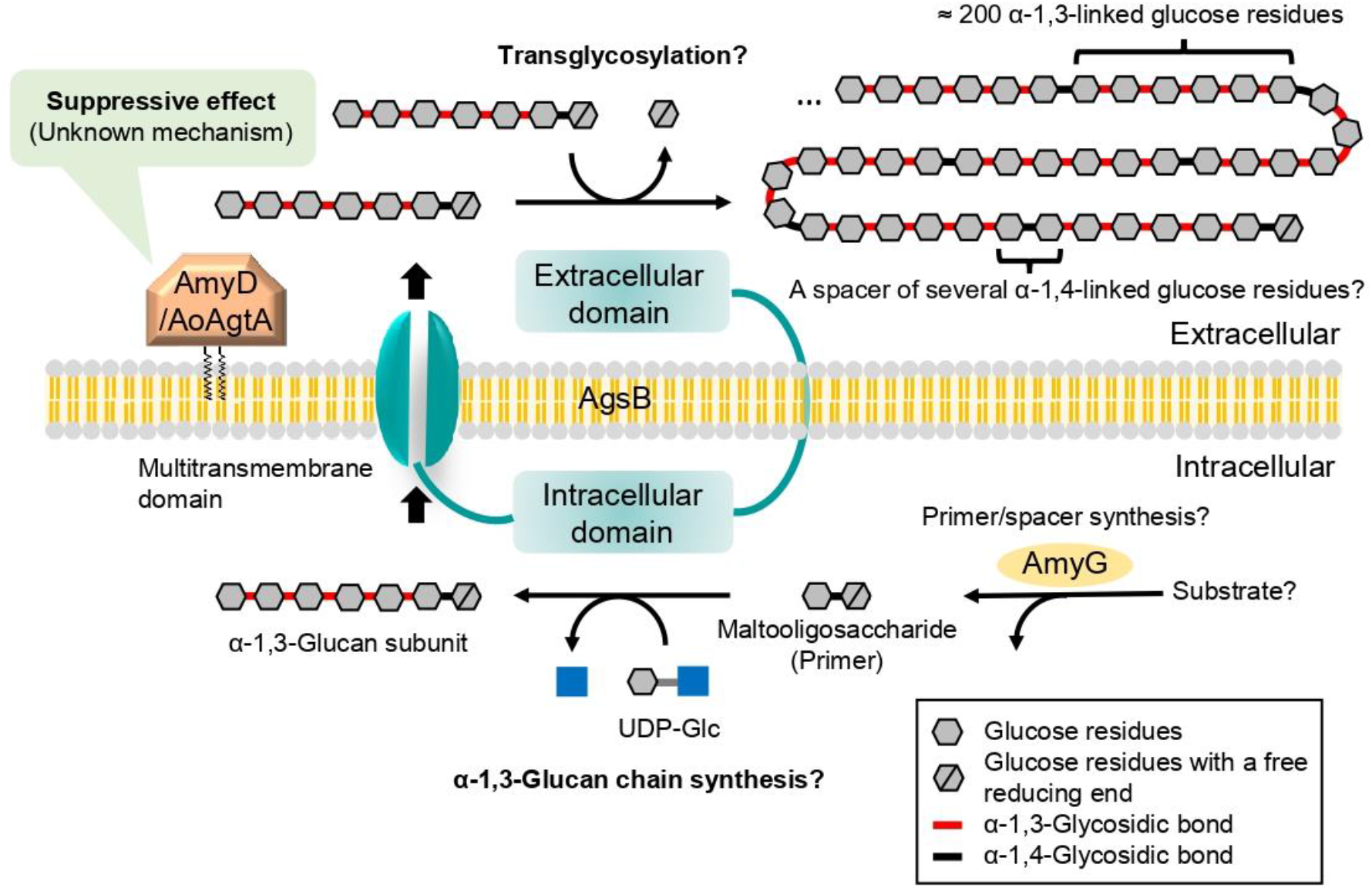
Speculative model for biosynthesis of cell wall α-1,3-glucan in *A. nidulans* and *A. oryzae*. Cell wall α-1,3-glucan is mainly synthesized by AgsB, which is composed of extracellular, intracellular, and multitransmembrane domains. Maltooligosaccharide is likely required as a primer for α-1,3-glucan biosynthesis and may be produced by intracellular α-amylase AmyG. The α-1,3-glucan chains may be synthesized by the intracellular domain of AgsB, and the resulting α-1,3-glucan units may be exported extracellularly by the multitransmembrane domain. The α-1,3-glucan units may be linked by transglycosylation catalyzed by the extracellular domain of AgsB to complete α-1,3-glucan maturation. Glycosylphosphatidylinositol-anchored α-amylase AmyD/AoAgtA regulates cell wall α-1,3-glucan biosynthesis, but the underlying mechanism remains unclear. Question marks indicate steps that are not firmly established.

Chemical analyses of cell wall α-1,3-glucan in *A. nidulans* (Miyazawa et al., 2018) and *Aspergillus wentii* (Choma et al., 2013) revealed concatenation of a subunit consisting of about 200 α-1,3-linked glucose residues and a spacer of several 1,4-linked glucose residues. The anomer of the glucose residues of the spacer in *Aspergillus* fungi has not been analyzed, but it is presumed to be α-type (Latgé, 2010).

AgtA of *Aspergillus oryzae* (AoAgtA) and proteins that are encoded by the orthologues of *A. oryzae agtA*, which are defined here as “Agt proteins”, are expected to be α-amylases of the glycoside hydrolase family 13 (GH13) and contain a C-terminal GPI-anchor site. He et al. (2014) reported that overexpression of *A. nidulans amyD*, orthologous to *A. oryzae agtA*, decreases the amount of cell wall α-1,3-glucan, whereas *amyD* disruption increases it, suggesting that *amyD* suppresses α-1,3-glucan biosynthesis. He et al. (2017) showed that the amount of cell wall α-1,3-glucan was lower in the *amyD*-overexpressing strain than in the parental *A. nidulans* strain at all time points examined, whereas in the α-1,3-glucanase overexpressing strains it was similar to that in the parental strain at the beginning of the time course and became lower from the middle of the culture. These results suggest that the mechanism of the decrease in the amount of cell wall α-1,3-glucan by AmyD differs from that by α-1,3-glucanase. We recently performed *in vivo* functional analysis of *amyD* in *A. nidulans* and found that not only the amount but also the molecular weight (MW) of cell wall α-1,3-glucan was decreased by *amyD* overexpression (Miyazawa et al., 2022). *Aspergillus niger* AgtA (AnAgtA) cannot use α-1,3-glucan derived from *A. nidulans* as a substrate and shows remarkable transglycosylation activity to produce maltooligosaccharides with a degree of polymerization (DP) of at least 28 (van der Kaaij et al., 2007). Overall, the mechanisms underlying the decrease in the amount and MW of α-1,3-glucan by Agt proteins are still unknown.

We have been developing *A. oryzae* strains with dispersed hyphae and high levels of enzyme production, which lack α-1,3-glucan and extracellular matrix galactosaminogalactan (Miyazawa et al., 2016, 2019; Ichikawa et al., 2022). Understanding the mechanism how AoAgtA regulates α-1,3-glucan biosynthesis would contribute to further development of phenotype control of *A. oryzae* from the viewpoint of hyphal pellet formation. There are three α-1,3-glucan synthase genes in *A. oryzae* (*agsA, agsB, agsC*), and *agsB* deletion leads to the loss of cell wall α-1,3-glucan (Zhang et al., 2017).

In the present study, we characterized an *agtA*-overexpressing (*agtA*^*OE*^) strain of *A. oryzae* and analyzed the enzymatic properties of recombinant (r)AoAgtA for maltooligosaccharides and their derivatives. By *in vivo* analysis of *A. nidulans* strains heterologously overexpressing *agtA*, we showed that AoAgtA decreases the MW of cell wall α-1,3-glucan. We discussed the enzymatic properties and the involvement of AoAgtA in cell wall α-1,3-glucan biosynthesis.

## MATERIALS AND METHODS

### Bioinformatics Tools

The signal peptide sequence was predicted by SignalP-5.0 Server (https://services.healthtech.dtu.dk/service.php?SignalP-5.0). The GPI anchor site was predicted by GPI Modification Site Prediction (https://mendel.imp.ac.at/gpi/gpi_server.html). The structure of AoAgtA was predicted using AlphaFold2 (Jumper et al., 2021; Tunyasuvunakool et al., 2021).

### Materials

*p-*Nitrophenyl α-maltopentaoside (Mal_5_-α-*p*NP) was synthesized in our laboratory (Usui and Murata, 1988). Maltooligosaccharides were kindly supplied by Kikkoman (Noda, Japan). Nigerooligosaccharides were prepared by two methods: (i) partial acid degradation of linear α-1,3-glucan produced by GtfJ, a glucosyltransferase from *Streptococcus salivarius* ATCC 25975 (Puanglek et al., 2016), according to Czerwonka et al. (2019); (ii) enzymatic synthesis using α-1,3-glucoside phosphorylase (Nihira et al., 2014). Dextran from *Leuconostoc mesenteroides* (average MW 9,000–11,000) was purchased from Merck (Darmstadt, Germany). Pullulan was kindly supplied by Dr. Tasuku Nakajima (Tohoku University) and nigeran by Dr. Keiko Uechi (University of the Ryukyus). Pustulan was purchased from Calbiochem (San Diego, CA, United States) and treated with alcohol as in Hattori et al. (2013). Laminaran from *Eisenia bicyclis* was purchased from Tokyo Chemical Industry (Chuo, Japan). All other commonly used chemicals were obtained from commercial sources.

### Strains

The *A. oryzae* and *A. nidulans* strains are listed in Table 1. *Escherichia coli* DH5α was used for plasmid amplification. *Pichia pastoris* SMD1168H (Thermo Fisher Scientific, Waltham, MA, United States) was used for protein expression.

**Table 1.**
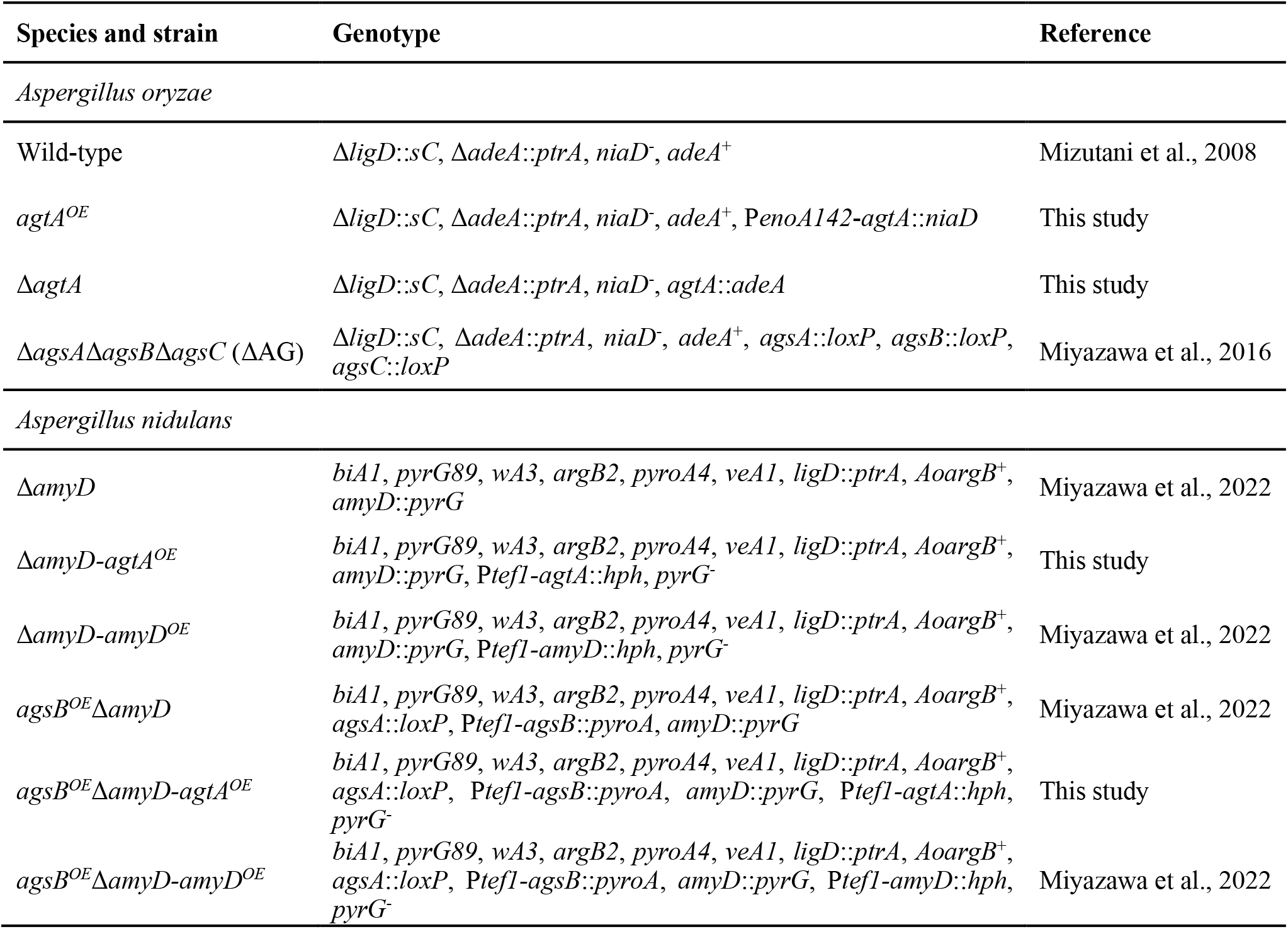
*Aspergillus* strains used in this study.

| Species and strain | Genotype | Reference |
| --- | --- | --- |
| *Aspergillus oryzae* |  |  |
| Wild-type | $\Delta ligD::sC$, $\Delta adeA::ptrA$, $niaD^-$, $adeA^+$ | Mizutani et al., 2008 |
| $agtA^{OE}$ | $\Delta ligD::sC$, $\Delta adeA::ptrA$, $niaD^-$, $adeA^+$, $P_{enoA142}-agtA::niaD$ | This study |
| $\Delta agtA$ | $\Delta ligD::sC$, $\Delta adeA::ptrA$, $niaD^-$, $agtA::adeA$ | This study |
| $\Delta agsA\Delta agsB\Delta agsC$ ($\Delta AG$) | $\Delta ligD::sC$, $\Delta adeA::ptrA$, $niaD^-$, $adeA^+$, $agsA::loxP$, $agsB::loxP$, $agsC::loxP$ | Miyazawa et al., 2016 |
| *Aspergillus nidulans* |  |  |
| $\Delta amyD$ | $biA1$, $pyrG89$, $wA3$, $argB2$, $pyroA4$, $veA1$, $ligD::ptrA$, $AoargB^+$, $amyD::pyrG$ | Miyazawa et al., 2022 |
| $\Delta amyD-agtA^{OE}$ | $biA1$, $pyrG89$, $wA3$, $argB2$, $pyroA4$, $veA1$, $ligD::ptrA$, $AoargB^+$, $amyD::pyrG$, $P_{tef1}-agtA::hph$, $pyrG^-$ | This study |
| $\Delta amyD-amyD^{OE}$ | $biA1$, $pyrG89$, $wA3$, $argB2$, $pyroA4$, $veA1$, $ligD::ptrA$, $AoargB^+$, $amyD::pyrG$, $P_{tef1}-amyD::hph$, $pyrG^-$ | Miyazawa et al., 2022 |
| $agsB^{OE}\Delta amyD$ | $biA1$, $pyrG89$, $wA3$, $argB2$, $pyroA4$, $veA1$, $ligD::ptrA$, $AoargB^+$, $agsA::loxP$, $P_{tef1}-agsB::pyroA$, $amyD::pyrG$ | Miyazawa et al., 2022 |
| $agsB^{OE}\Delta amyD-agtA^{OE}$ | $biA1$, $pyrG89$, $wA3$, $argB2$, $pyroA4$, $veA1$, $ligD::ptrA$, $AoargB^+$, $agsA::loxP$, $P_{tef1}-agsB::pyroA$, $amyD::pyrG$, $P_{tef1}-agtA::hph$, $pyrG^-$ | This study |
| $agsB^{OE}\Delta amyD-amyD^{OE}$ | $biA1$, $pyrG89$, $wA3$, $argB2$, $pyroA4$, $veA1$, $ligD::ptrA$, $AoargB^+$, $agsA::loxP$, $P_{tef1}-agsB::pyroA$, $amyD::pyrG$, $P_{tef1}-amyD::hph$, $pyrG^-$ | Miyazawa et al., 2022 |

### Construction of an *agtA*-Overexpressing Strain and an *agtA* Gene Disruptant in *Aspergillus oryzae*

Primers are listed in Supplementary Table 1. An *agtA* overexpression plasmid, pNEN142-agtA, was constructed as follows. The *agtA* gene was amplified by polymerase chain reaction (PCR) with the agtA-Fw-NotI and agtA-Rv-NotI primers (designed with a NotI restriction site each) and wild-type *A. oryzae* genomic deoxyribonucleic acid (DNA) as a template (Supplementary Figure 1A). The PCR product was ligated into the NotI site of the pNEN142 vector (Minetoki et al., 2003) with an In-Fusion HD Cloning Kit (Takara Bio, Kusatsu, Japan); this vector contains the improved promoter of the *A. oryzae enoA* gene. The pNEN142-agtA plasmid was transformed into wild-type *A. oryzae* (Supplementary Figure 1B). The *agtA*^*OE*^ transformants were selected on standard Czapek–Dox (CD) medium (Miyazawa et al., 2019); the *niaD* gene was used as a selectable marker. The transformants were subjected to single sporing as described previously (Hara et al., 2002). Integration of the *agtA* overexpression cassette was confirmed by PCR (Supplementary Figure 1C).

The *agtA*-disruption (Δ*agtA*) strain was created as described previously (Tamano et al., 2007). The *agtA* gene disruption cassette was generated by fusion PCR using an Expand High Fidelity PCR System (F. Hoffmann-La Roche, Basel, Switzerland). The 5′- and 3′-arms of *agtA* were amplified from wild-type *A. oryzae* genomic DNA with the primers agtA-LU and agtA-LL+adeA for the 5′-arm, and agtA-RU+adeA and agtA-RL for the 3′-arm (Supplementary Figure 2A. *adeA* was amplified with the primer pair agtA-AU and agtA-AL (Supplementary Figure 2A). A mixture of the 5′-flanking amplicon: *adeA*: 3′-flanking amplicon at a 1: 3: 1 molar ratio was used as a template. The PCR products were used for a second PCR round with the primer pair agtA-LU and agtA-RL to fuse the 5′ and 3′ regions of the target gene at each end of the *adeA* gene (Supplementary Figure 2A). The amplified fragment was transformed into wild-type *A. oryzae* (Supplementary Figure 2B). The transformants were subjected to two consecutive rounds of single sporing. Replacement of the *agtA* gene was confirmed by PCR (Supplementary Figure 2C).

### Analysis of Growth Characteristics of *Aspergillus oryzae* and *Aspergillus nidulans* in Submerged Culture

Conidia (final concentration, 1 × 10^5^ /mL) of *A. oryzae* strains were inoculated into 50 mL of YPD medium containing 1% yeast extract, 2% peptone, and 2% glucose in 200-mL Erlenmeyer flasks and rotated at 120 rpm at 30°C for 24 h. The mean diameter of the hyphal pellets was determined as described previously (Miyazawa et al., 2016). Conidia (final concentration, 5 × 10^5^ /mL) of *A. nidulans* strains were inoculated into 50 mL of CD medium (Miyazawa et al., 2018) in 200-mL Erlenmeyer flasks and rotated at 160 rpm at 37°C for 24 h.

### α-1,3-Glucan Quantification in *Aspergillus oryzae* and *Aspergillus nidulans* Mycelia

Strains of *A. oryzae* were inoculated into YPD medium and cultured as described in the Analysis of Growth Characteristics subsection. Strains of *A. nidulans* were inoculated into 200 mL of CD medium in 500-mL Erlenmeyer flasks and rotated at 160 rpm at 37°C for 24 h. Mycelia were collected by filtration through Miracloth (Merck), washed with distilled water, lyophilized, and pulverized in a Mixer Mill MM 400 (Retsch, Haan, Germany). The resulting *A. nidulans* powder (0.5 g) was suspended in chloroform–methanol (3:1 vol/vol) and delipidized as described previously (Miyazawa et al., 2022). The pulverized *A. oryzae* mycelia (0.25–1 g) or delipidized *A. nidulans* mycelia were suspended in 0.1 M sodium phosphate buffer (pH 7.0). Cell wall components were fractionated by hot-water and alkali treatment (Yoshimi et al., 2013); the fractionation resulted in hot-water-soluble, alkali-soluble (AS), and alkali-insoluble fractions. The AS fraction was further separated into a fraction soluble in water at neutral pH (AS1) and an insoluble fraction (AS2). The AS2 fraction was used as α-1,3-glucan; this fraction was hydrolyzed and its glucose content was quantified as described previously (Yoshimi et al., 2013).

### Construction of a Recombinant AoAgtA Expression Strain in *Pichia pastoris*

An *agtA*-expression plasmid, pPICZα B-agtA, was constructed as follows. The gene encoding AoAgtA without the C-terminus (amino acids (aa) 1–518; whole protein, 549 aa) to prevent GPI-anchoring was amplified with the primers agtA-Fw-NdeI (designed with a NdeI restriction site) and agtA-Rv-SmaI (designed with a SmaI restriction site) from wild-type *A. oryzae* complementary DNA (cDNA) as a template. The PCR product was digested with NdeI and SmaI and ligated into the NdeI– SmaI site of the pIVEX 2.3d vector (F. Hoffmann-La Roche). Then the AoAgtA coding region without the 1st–23th aa (putative signal peptide) and with a PGGGS linker and a (His)_6_ tag at the C-terminus was amplified with the primers agtA-Fw-PstI (designed with a PstI restriction site) and agtA-Rv-XbaI (designed with a XbaI restriction site); pIVEX 2.3d-agtA was used as a template. The PCR product was digested with PstI and XbaI and ligated into the PstI–XbaI site of the pPICZα B vector (Thermo Fisher Scientific), which encodes a secretion signal of *Saccharomyces cerevisiae* α-factor. The pPICZα B-agtA plasmid was linearized by SacI and integrated into the chromosomal DNA of *P. pastoris* SMD1168H competent cells with an EasySelect *Pichia* Expression Kit (Thermo Fisher Scientific) according to the manufacturer’s instructions. The *agtA-*expressing transformants were cultured on YPDS (1% yeast extract, 2% peptone, 2% glucose, and 1 M sorbitol) agar plates containing zeocin (100 µg/mL) at 30°C for 2 days.

### Expression and Purification of Recombinant AoAgtA

Buffered glycerol-complex (BMGY) medium and buffered methanol-complex (BMMY) medium were prepared according to the instructions of the EasySelect *Pichia* Expression Kit. The rAoAgtA-expressing *P. pastoris* strain was inoculated into 25 mL of BMGY medium in a 100-mL Erlenmeyer flask and rotated at 160 rpm at 30°C until OD_600_ 4. The culture broth was centrifuged at 3,000 × *g* for 5 min. The collected cells were resuspended into 100 mL BMMY medium in a 500-mL Erlenmeyer flask and rotated at 200 rpm at 30°C for 6 days. To induce and maintain the expression of rAoAgtA, methanol (final concentration, 0.5% vol/vol) was added to culture broth every 24 h. The culture broth was centrifuged at 3,000 × *g* for 5 min, and the supernatant was dialyzed against 10 mM Tris-HCl buffer (pH 8.0). The supernatant (25 mL) was applied to a Ni Sepharose 6 Fast Flow column (1.8 × 5 cm; Cytiva, Marlborough, MA, United States) equilibrated in 20 mM Tris-HCl buffer (pH 8.0). The column was washed with the same buffer, and bound enzyme was eluted with 250 mM imidazole in the same buffer with pH readjusted to 8.0. The eluant was concentrated and buffer was replaced with 10 mM Tris-HCl buffer (pH 8.0) in Amicon Ultra-15 (nominal MW limit, 10,000) (Merck). At each stage, protein content was determined by Bradford method with bovine serum albumin as a standard, and rAoAgtA activity was determined as in the Recombinant AoAgtA Activity Measurement subsection. To check protein purity and determine the MW of the purified protein, it was subjected to sodium dodecylsulfate-polyacrylamide gel electrophoresis (SDS-PAGE) (Laemmli, 1970), and the gels were stained with Coomassie Brilliant Blue R-250; the MW standards (10,000–250,000) were Precision Plus Protein Kaleidoscope Standards (Bio-Rad Laboratories, Hercules, CA, United States). Purified protein (4 μg) was deglycosylated with 500 U of Endoglycosidase H (Endo H) (New England Biolabs, Ipswich, MA, United States) according to the manufacturer’s instructions.

### Analytical High-Performance Liquid Chromatography

The high-performance liquid chromatography (HPLC) analysis was carried out with a Jasco Intelligent System Liquid Chromatograph (Jasco, Hachioji, Japan) under Conditions 1 and 2. Under Condition 1, a Unison UK-C18 column (4.6 × 250 mm; Imtakt, Kyoto, Japan) was used, and the detection was performed at 300 nm. The bound material was eluted with 20% methanol at a flow rate of 1.0 mL/min at 40°C. Under Condition 2, a Shodex HILICpak VG-50 4E column (4.6 × 250 mm; Showa Denko, Minato, Japan) was used, and the detection was performed with a refractive index detector. The bound material was eluted with 65% acetonitrile at a flow rate of 0.7 mL/min at 40°C. Under Condition 3, the HPLC analysis was carried out with a Hitachi Elite LaChrom HPLC System (Hitachi, Chiyoda, Japan). The same Shodex column as under Condition 2 was used, and the detection was performed with an evaporative light scattering detector (Alltech 3300 ELSD; Buchi Labortechnik, Flawil, Switzerland). The bound material was eluted with 65% acetonitrile at a flow rate of 0.5 mL/min at 40°C. When the products of substrate degradation by rAoAgtA were analyzed by HPLC, data analysis was performed as follows. Products formed from the initial substrate at an early stage (up to approximately 15% degradation) were analyzed. Under Condition 1, the amount of each *p*-nitrophenylated product was calculated by multiplying the ratio of each peak area to the total peak area in the chromatogram by the concentration of the initial substrate. Under Conditions 2 and 3, the amount of each product was calculated from each peak area on the chromatogram and the calibration curve for each oligosaccharide.

### Recombinant AoAgtA Activity Measurement

A mixture (20 μL) containing 1 mM Mal_5_-α-*p*NP and an appropriate amount of rAoAgtA in 50 mM sodium acetate (Na-Ac) buffer (pH 5.5) was incubated at 40°C for 10 min. Aliquots (2 μL) were taken at 2-min intervals, and the reaction was immediately stopped with 40 μL of methanol. Then H_2_O (158 µL) was added to each aliquot, and the samples were subjected to HPLC analysis under Condition 1. The release velocity of *p*-nitrophenyl α-maltoside (Mal_2_-α-*p*NP) was determined as the slope of the time-course plot of Mal_2_-α-*p*NP amount. One unit of rAoAgtA activity was defined as the amount of enzyme required to liberate 1 μmol of Mal_2_-α-*p*NP from Mal_5_-α-*p*NP per minute.

### Substrate Specificity of Recombinant AoAgtA

Corn starch, potato starch, soluble starch, dextran, pullulan, bacterial α-1,3-glucan, nigeran, cellulose, pustulan, and laminaran were tested as rAoAgtA substrates. A mixture (12 μL) containing 2 mg/mL (0.2%) each substrate and 24 mU/mL purified rAoAgtA in 50 mM Na-Ac buffer (pH 5.5) was incubated at 40°C for 4 h. Then, H_2_O (138 μL) was added, and the reaction was immediately stopped by boiling for 5 min. The amount of reducing sugar generated from each substrate was measured by bicinchoninic acid method (Doner and Irwin, 1992; Utsumi et al., 2009). The bicinchoninic acid working reagent was prepared according to Utsumi et al. (2009), and 100 μL was added to a 100-μL aliquot of each boiled reaction mixture. The sample was incubated at 80°C for 40 min and then at room temperature for 15 min, and the absorbance at 560 nm was measured with a Multiscan Spectrum spectrophotometer (Thermo Fisher Scientific). One unit of enzyme activity was defined as the amount of enzyme required to liberate 1 μmol of reducing sugars (D-glucose conversion) from the substrate per minute. The detection limit was 1 mU/mL or less.

### Analysis of the Modes of Bond Cleavage in Maltooligosaccharides and Their *p*-Nitrophenyl Derivatives

Maltooligosaccharides (DP 2–8, Mal_2–8_) and *p-*nitrophenyl α-maltooligosides (PNMs, DP 2–8, Mal_2– 8_-α-*p*NP) were used to determine the modes of enzymatic bond cleavage by rAoAgtA. PNMs were synthesized from Mal_5_-α-*p*NP by transglycosylation catalyzed by rAoAgtA (Supplementary Material; Supplementary Figure 3). To test maltooligosaccharides, a mixture (40 μL) containing 25 mM each substrate and 9.5 mU/mL purified rAoAgtA in 50 mM Na-Ac buffer (pH 5.5) was incubated at 40°C for 20 min. An aliquot (10 μL) was taken, and the reaction was immediately stopped with 65 μL of acetonitrile. Then H_2_O (25 µL) was added to the aliquot, and the sample was subjected to HPLC analysis under Condition 2. The frequency of rAoAgtA-catalyzed cleavages of glycosidic linkages was calculated from the amount of each product.

To test PNMs, a mixture (20 μL) containing 1.6 mM each substrate and 9.5 mU/mL purified rAoAgtA in 50 mM Na-Ac buffer (pH 5.5) was incubated at 40°C for 5 min. Aliquots (2 μL) were taken at 1-min intervals, and the reaction was immediately stopped with 40 μL of methanol. Then H_2_O (158 µL) was added to each aliquot, and the samples were subjected to HPLC analysis under Condition 1. The release velocity of each PNM from the initial substrate was determined as the slope of the time-course plot of the amount of that PNM. The frequency of rAoAgtA-catalyzed cleavages of glycosidic linkages was calculated from the release velocities of different PNMs.

### Kinetic Studies

Seven substrate concentrations ([S]) for Mal_5_-α-*p*NP (1.6–100 mM), six each for Mal_6_-α-*p*NP and Mal_7_-α-*p*NP (1.6–50 mM), and five for Mal_8_-α-*p*NP (1.6–25 mM) were used. The degradation reactions of these PNMs, including hydrolysis and transglycosylation, catalyzed by rAoAgtA and HPLC analyses were performed as described in the Analysis of the Modes of Bond Cleavage subsection. The initial velocity (*v*) of substrate degradation was determined as the slope of the time-course plot of the total amount of all PNMs liberated from each initial substrate. The kinetic parameters of the Michaelis–Menten equation were evaluated by Hanes–Woolf plots ([S]/*v versus* [S]) and the least-squares method. The initial concentration of rAoAgtA in the reaction solution was 1.83 × 10^−4^ mM.

### Effects of Nigerooligosaccharides on Recombinant AoAgtA Catalytic Activity

Mal_5_-α-*p*NP and various sugars (nigerose, nigerotriose, and glucose) were prepared at a final concentration of 1.6 mM and 16 mM, respectively. The reactions and HPLC analyses were performed as described in the Analysis of the Modes of Bond Cleavage subsection, and the release velocity of Mal_2_-α-*p*NP was calculated.

### Enzymatic Synthesis of 3-α-Maltooligosylglucose

A mixture (530 μL) containing Mal_5_-α-*p*NP (50 mg, 100 mM), nigerose (100 mg, 560 mM), and 180 mU/mL rAoAgtA in dialyzed culture supernatant in 50 mM Na-Ac buffer (pH 5.5) was incubated at 40°C for 24 h. The reaction was stopped by adding 11 mL of methanol. The reaction mixture was dried, dissolved in a small amount of H_2_O, and then applied to an ODS column (1.3 × 50 cm; Yamazen, Osaka, Japan) equilibrated with H_2_O at a flow rate of 3.0 mL/min. The eluate (800 mL) was collected and dried. The partially purified products were dissolved in a small amount of H_2_O, and then applied to a Toyopearl HW-40S column (3.5 × 62 cm; Tosoh, Minato, Japan) equilibrated with H_2_O at a flow rate of 0.5 mL/min. The eluate was collected in 2-mL fractions (300 mL in total). Each fraction was analyzed by HPLC under Condition 3, and the fractions containing 3-α-maltotetraosylglucose (Mal_4_α1,3Glc), 3-α-maltotriosylglucose (Mal_3_α1,3Glc), 3-α-maltosylglucose (Mal_2_α1,3Glc), and nigerose were concentrated and then lyophilized. Fractions containing products of insufficient purification were rechromatographed under the same conditions. Finally, Mal_4_α1,3Glc (7.3 mg, yield 16.7% based on Mal_5_-α-*p*NP), Mal_3_α1,3Glc (3.1 mg, 8.8%), Mal_2_α1,3Glc (10.8 mg, 40.7%), and nigerose (65.7 mg) were obtained.

The structures of the synthesized products were evaluated by ^1^H and ^13^C nuclear magnetic resonance (NMR) analysis in D_2_O (Supplementary Tables 2, 3); 500 MHz ^1^H NMR spectra and 125 MHz ^13^C NMR spectra were recorded using a Bruker Avance Neo-500 NMR spectrometer (Bruker, Billerica, MA, United States). Matrix assisted laser desorption/ionization-time of flight (MALDI-TOF) mass spectra were acquired using an Autoflex Speed spectrometer (Bruker). MALDI-TOF mass analysis of 3-α-maltooligosylglucose showed *m*/*z* 527.167 [M + Na]^+^ (calcd for C_18_H_32_NaO_16_, 527.159), 689.210 [M + Na]^+^ (calcd for C_24_H_42_NaO_21_, 689.212), and 851.392 [M + Na]^+^ (calcd for C_30_H_52_NaO_26_, 851.264).

### Behavior Analysis of Recombinant AoAgtA for 3-α-Maltotetraosylglucose

A mixture (12 μL) containing 25 mM Mal_4_α1,3Glc and 180 mU/mL purified rAoAgtA in 50 mM Na-Ac buffer (pH 5.5) was incubated at 40°C for 30 min. An aliquot (3 µL) was taken, and the reaction was immediately stopped with 44 µL of acetonitrile. Then H_2_O (21 µL) was added to the aliquot, and the sample was subjected to HPLC analysis under Condition 3. The substrate-degradation velocity and the frequency of rAoAgtA-catalyzed cleavages of glycosidic linkages were determined from the amounts of degradation products. The substrates Mal_5_ and nigeropentaose were used for comparison. For Mal_5_, the reaction time was set to 8 min and degradation velocity was determined from the amount of Mal_2_ liberated.

### Construction of *agtA*-Overexpressing Strains in *Aspergillus nidulans*

The *agtA*^*OE*^ strains were constructed by inserting the *agtA* overexpression cassette into the disrupted *amyD* locus. The pNEN142-agtA(-intron) and pAHdPT-agtA plasmids were first constructed (Supplementary Figure 4A). To remove the intron in the open reading frame of *agtA*, PCR was performed using PrimeSTAR Max DNA Polymerase (Takara Bio) and the pNEN142-agtA plasmid as a template. The product was transformed into *E. coli* to obtain the pNEN142-agtA(-intron) plasmid. To construct pAHdPT-agtA, a fragment containing the *agtA* open reading frame and *agdA* terminator was amplified from pNEN142-agtA(-intron). Primers AopyrG-IF-Right-Fw and Ptef1-tail-Rv were used in PCR with pAHdPT-amyD (Miyazawa et al., 2022) as a template. The two fragments were fused using the In-Fusion HD Cloning Kit according to the manufacturer’s instructions. The resulting plasmid pAHdPT-agtA was digested with SacI and transformed into the Δ*amyD* and *agsB*^*OE*^Δ*amyD* strains (Supplementary Figure 4B). Candidate strains were selected on CD medium containing uridine, uracil, and 1.3 mg/mL 5-fluoroorotic acid and then cultured on CD medium containing uridine, uracil, and 800 µg/mL hygromycin. Correct insertion of the cassette was confirmed by PCR (Supplementary Figure 4C).

### Determination of the Average Molecular Weight of Alkali-Soluble Glucan

The MW of glucan in the AS2 fraction was determined by gel permeation chromatography according to Miyazawa et al. (2022). Polystyrene (MW, 13,900–3,850,000; Showa Denko) was used as a standard to calibrate the column.

## RESULTS

### Sequence Analysis of AoAgtA

According to the NCBI database, the *agtA* (AO090003001498) gene in *A. oryzae* consists of 1,711 base pairs and encodes a 549-aa protein with a putative MW of 60,400. A comparison of the cDNA and genome sequence revealed one intron. The predicted signal peptide (1–23 aa) and a GPI anchor site (ω site, Ser 530) suggested that AoAgtA is a cell membrane and/more cell wall-bound protein.

The aa sequence of AoAgtA showed 44% identity with that of Taka-amylase A of *A. oryzae* (encoded by *amyB* and *amyC*; hereinafter TAA) belonging to GH13. A comparison between the AlphaFold2-predicted model of AoAgtA and the TAA crystal structure (PDB, 2GVY) (Vujičić-žagar and Dijkstra, 2006) showed that the overall structure, catalytic residues, and Ca^2+^-binding residues were highly conserved (Supplementary Figure 5).

### The *agtA*-Overexpressing Strain of *Aspergillus oryzae*

We constructed the *agtA*^*OE*^ and Δ*agtA* strains from wild-type *A. oryzae* (Supplementary Figures 1, 2). In submerged culture, the size of the hyphal pellets of the *agtA*^*OE*^ strain was as small as that of the Δ*agsA*Δ*agsB*Δ*agsC* (ΔAG) strain lacking cell wall α-1,3-glucan (Figures 2A, B). We quantified the amount of glucose in the AS2 fraction obtained from lyophilized mycelia of each strain; this fraction contains mainly α-1,3-glucan. The proportion of glucose in the AS2 fraction from the *agtA*^*OE*^ strain was reduced to 24% of that from the wild-type (Figure 2C; *P* < 0.01), whereas the size of hyphal pellets and the α-1,3-glucan content of Δ*agtA* strain were similar to those of the wild-type (Figure 2). These results suggest that AoAgtA negatively regulates cell wall α-1,3-glucan biosynthesis in *A. oryzae*.

**Figure 2.**
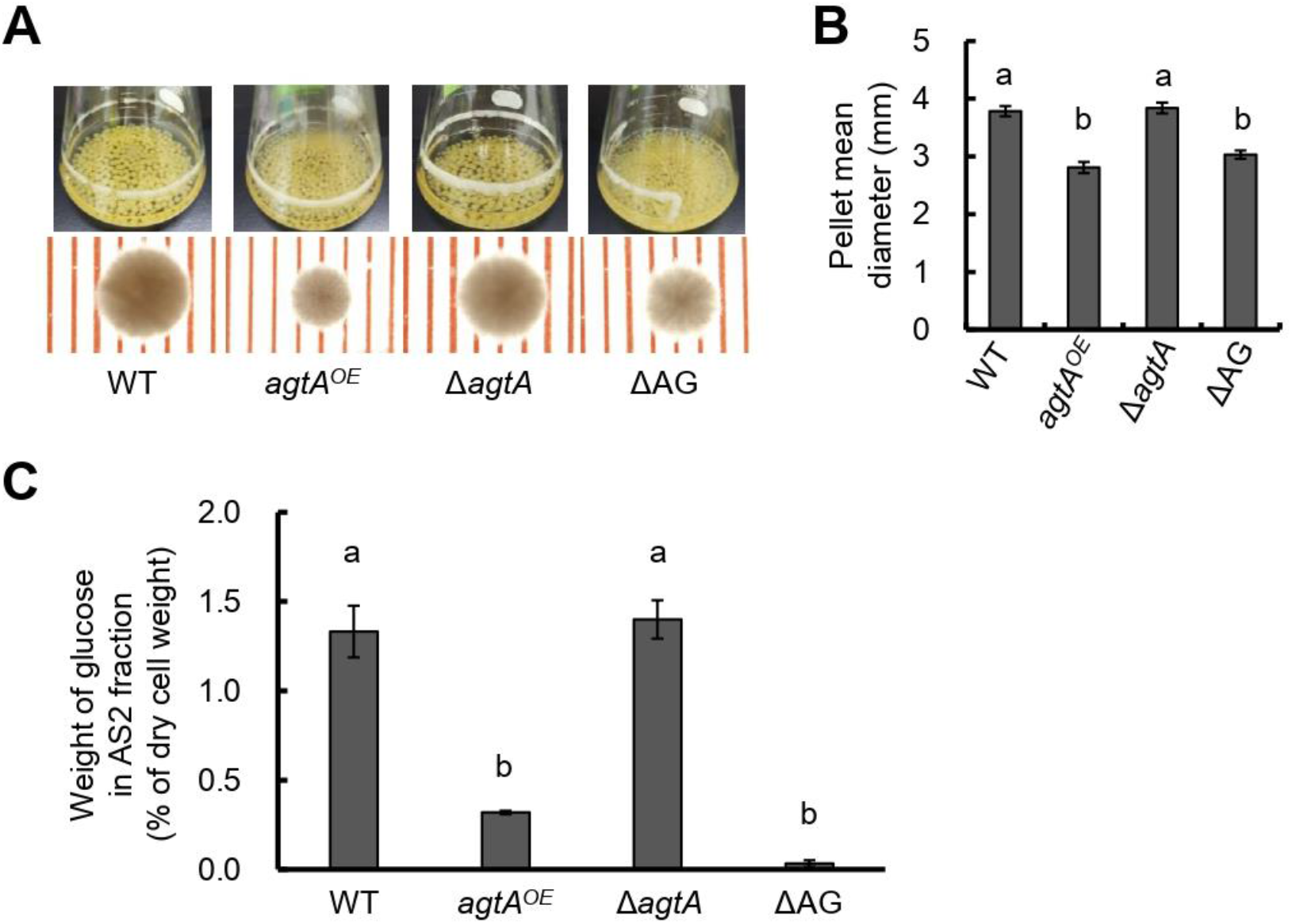
Analysis of the *agtA*^*OE*^ strain of *A. oryzae*. **(A)** Growth characteristics of *A. oryzae* wild-type (WT), *agtA*^*OE*^, Δ*agtA*, and ΔAG strains in submerged culture. Conidia (final concentration, 1 × 10^5^ /mL) of each strain were inoculated into 50 mL of YPD medium and rotated at 120 rpm at 30°C for 24 h. Upper images, cultures in Erlenmeyer flasks; lower images, representative hyphal pellets under a stereomicroscope (scale = 1 mm). **(B)** The mean diameter of hyphal pellets was determined by measuring 10 randomly selected pellets under a stereomicroscope. Error bars represent the standard error of the mean calculated from three replicates. Different letters above bars indicate significant difference by Tukey’s test (*P* < 0.01). **(C)** Amount of glucose in the alkali-soluble water-insoluble (AS2) fraction. Error bars represent the standard error of the mean calculated from three replicates. Different letters above bars indicate significant difference by Tukey’s test (*P* < 0.01).

### Recombinant AoAgtA Production in *Pichia pastoris*

Because *agtA* overexpression in *A. oryzae* led to a decrease in the amount of cell wall α-1,3-glucan and consequently to the formation of small hyphal pellets, we purified and characterized rAoAgtA produced in *P. pastoris*. In a single-step purification of (His)_6_-tagged rAoAgtA, 22% of enzymatic activity was recovered from culture supernatant on day 6 of culture; 5.2-fold purification was achieved (Table 2). Purified rAoAgtA showed a smeared band on SDS-PAGE with a MW of 73,200 (Figure 3). Removal of *N*-glycosylation by Endo H treatment decreased the apparent protein MW to 51,700 (Figure 3); the expected MW of rAoAgtA without the signal peptide and GPI anchor site but with the linker and the (His)_6_ tag at the C-terminus is 56,800. Therefore, the Endo H treatment revealed *N*-glycosylation of rAoAgtA.

**Figure 3.**
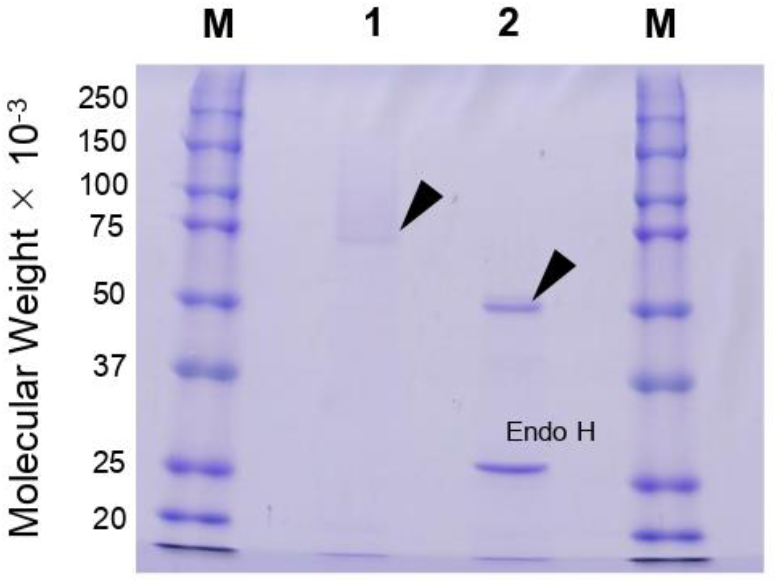
SDS-PAGE analysis of rAoAgtA produced in *P. pastoris*. Lanes M, Molecular weight markers; Lane 1, rAoAgtA purified on a Ni Sepharose 6 Fast Flow column; Lane 2, The same rAoAgtA deglycosylated with Endo H. Arrowheads indicate rAoAgtA bands. The band at 29,000 in lane 2 is Endo H.

**Table 2.** Purification of rAoAgtA from the culture supernatant of the rAoAgtA-expressing *P. pastoris* strain.

| Step | Total activity (U) | Protein (mg) | Specific activity (U/mg) | Yield (%) | Purification (fold) |
| --- | --- | --- | --- | --- | --- |
| Culture supernatant | 0.17 | 1.6 | 0.11 | 100 | 1 |
| Ni Sepharose 6 Fast Flow eluate | 0.039 | 0.069 | 0.56 | 22 | 5.2 |

### Biochemical Characterization of Recombinant AoAgtA

To characterize the biochemical properties of rAoAgtA, we investigated optimal conditions and stability as described in Supplementary Material. The optimal temperature of rAoAgtA was 40°C (Supplementary Figure 6A). After 30-min incubation at 10–70°C, the protein retained >80% of enzymatic activity (i.e., was stable) at up to 30°C (Supplementary Figure 6B). The optimal pH of rAoAgtA was 5.5 (Supplementary Figure 6C). After 30-min incubation in each buffer at 4°C, the protein retained >80% enzymatic activity (i.e., was stable) at pH 4–9 (Supplementary Figure 6D).

Cu^2+^ and ethylenediaminetetraacetic acid (EDTA) reduced rAoAgtA activity to undetectable and 21 ± 6%, respectively (Supplementary Table 4). rAoAgtA activity was restored by adding Ca^2+^ after EDTA treatment (Supplementary Table 5). The predicted conservation of the Ca^2+^-binding residues between AoAgtA and TAA (Supplementary Figure 5E) is consistent with these results.

### Substrate Specificity of Recombinant AoAgtA

We assessed the ability of purified rAoAgtA to catalyze the degradation of various natural glucans by measuring the amounts of reducing sugars produced from the substrates. rAoAgtA showed a weak degradation activity (3.23 ± 0.28 mU/mL) only for soluble starch among glucans (corn starch, potato starch, soluble starch, dextran, pullulan, and nigeran) containing α-1,4-glycosidic bonds (Supplementary Table 6). Furthermore, rAoAgtA was unable to degrade bacterial α-1,3-glucan that contained only α-1,3-glycosidic bonds, as well as cellulose, pustulan, and laminaran (Supplementary Table 6). Since rAoAgtA did not degrade bacterial α-1,3-glucan and nigeran, which also contains α-1,3-glycosidic bonds, AoAgtA appears not to cleave the α-1,3-glycosidic bonds in cell wall α-1,3-glucan.

### Analysis of the Modes of Bond Cleavage in Maltooligosaccharides and Their *p*-Nitrophenyl Derivatives

Since rAoAgtA degraded soluble starch, we evaluated its degradation activity and the modes of bond cleavage for 25 mM maltooligosaccharides (Mal_2–8_) and 1.6 mM their *p*NP-derivatives (Mal_2–8_-α-*p*NP). The cleavage positions and frequencies of glycosidic bonds degraded by rAoAgtA in each oligosaccharide substrate are shown in Table 3. rAoAgtA showed degradation activity against maltooligosaccharide substrates with a DP of at least 5 (Mal_5–8_ and Mal_5–8_-α-*p*NP). HPLC chromatograms of the substrates and products of Mal_5–8_-α-*p*NP degradation by rAoAgtA are shown in Supplementary Figure 7. rAoAgtA catalyzed not only hydrolysis but also transglycosylation. For example, when the substrate was Mal_5_, the degradation products Mal_2_ and Mal_3_, and the transglycosylation product Mal_8_ were observed. Since the molar concentration of Mal_2_ was equal to the sum of molar concentrations of Mal_3_ and Mal_8_ (data not shown), we concluded that the cleavage position in Mal_5_ was the third glycosidic bond from the non-reducing end (Table 3). In maltooligosaccharide substrates (DP of at least 5) other than Mal_5_ and Mal_5_-α-*p*NP, rAoAgtA randomly cleaved the internal α-1,4-glycosidic bonds (Table 3). The mode of bond cleavage by rAoAgtA in maltooligosaccharide substrates with a DP of at least 5 was similar to that of TAA, which is a typical α-amylase (Nitta et al., 1971; Suganuma, 1983). On the other hand, a major difference was observed in that TAA can hydrolyze Mal_2–4_ (Nitta et al., 1971; Suganuma, 1983), whereas rAoAgtA cannot.

**Table 3.**
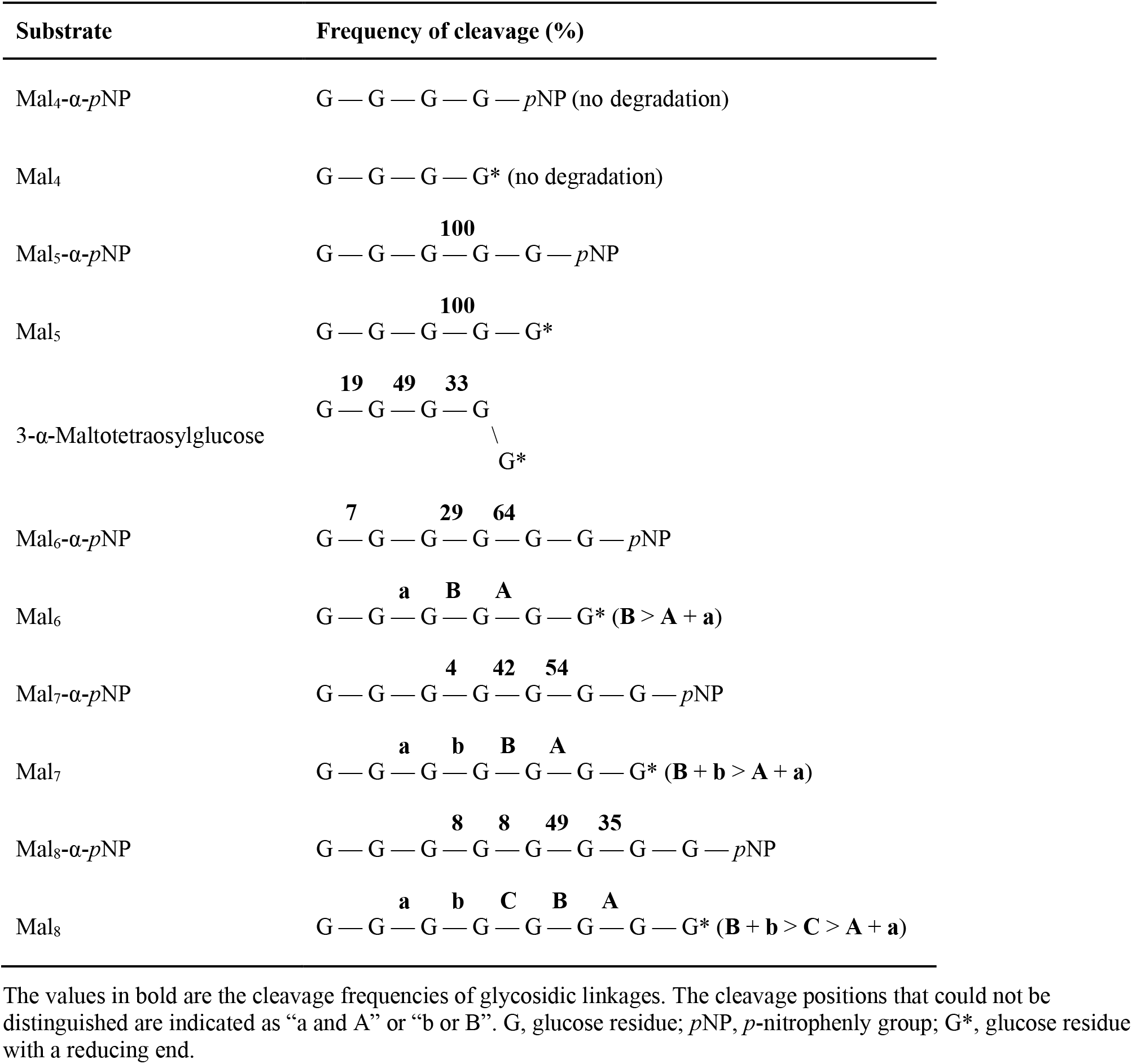
Frequency of site-specific cleavage during rAoAgtA-catalyzed degradation of maltooligosaccharides and their derivatives.

### Kinetic Studies

We conducted kinetic studies of the degradation of PNMs (Mal_5–8_-α-*p*NP) by rAoAgtA to further investigate its properties (Table 4, Supplementary Figure 8). Ideally, the kinetics should be analyzed in the absence of transglycosylation, but we could not prevent transglycosylation. Therefore, we analyzed the kinetics in the presence of PNM hydrolysis and transglycosylation to the initial substrate, and obtained the apparent kinetic parameter values. The *K*_m_ value of rAoAgtA was the lowest for Mal_8_-α-*p*NP (25.1 mM), and the *k*_cat_/*K*_m_ value was the highest for Mal_6_-α-*p*NP (1.42 s^-1^mM^-1^). Other known α-amylases often have *K*_m_ values of at most several mM and larger *k*_cat_/*K*_m_ values than those of rAoAgtA for maltooligosaccharide substrates with a DP at least 5 (Nitta et al., 1971; Suganuma, 1983; Usui et al., 1992; Okada et al., 2000). Thus, the affinity and catalytic efficiency of rAoAgtA for maltooligosaccharide substrates were lower than those of other known α-amylases.

**Table 4.** Kinetic parameters of rAoAgtA for the degradation of Mal_5–8_-α-*p*NP.

| Substrate | $K_m$ (mM) | $V_{max}$ (mM/min) | $k_{cat}$ (1/s) | $\frac{k_{cat}}{K_m}$<br>(1/(s · mM)) | Relative $k_{cat}/K_m$ |
| --- | --- | --- | --- | --- | --- |
| Mal<sub>5</sub>-α-pNP | 61.8 | 0.659 | 60.0 | 0.97 | 0.69 |
| Mal<sub>6</sub>-α-pNP | 45.6 | 0.710 | 64.6 | 1.42 | 1.00 |
| Mal<sub>7</sub>-α-pNP | 28.9 | 0.402 | 36.7 | 1.27 | 0.90 |
| Mal<sub>8</sub>-α-pNP | 25.1 | 0.344 | 31.3 | 1.25 | 0.88 |

The relative $k_{cat}/K_m$ values were calculated by dividing the $k_{cat}/K_m$ values by that for Mal<sub>6</sub>- $\alpha$ -pNP, which was the largest among those determined.

### Effects of Nigerooligosaccharides on Recombinant AoAgtA Catalytic Activity

Nigerooligosaccharides can be considered as part of the structure of cell wall α-1,3-glucan of *Aspergillus* fungi. In the presence of rAoAgtA, 1.6 mM Mal_5_-α-*p*NP as a substrate, and 16 mM nigerose or nigerotriose, the release velocity of Mal_2_-α-*p*NP was 1.2 times and 1.4 times, respectively, that in the control without nigerooligosaccharides (Supplementary Figure 9; *P* < 0.01). Addition of glucose instead of nigerooligosaccharides had no effect (Supplementary Figure 9). This may be a result of substrate degradation due to the occurrence of transglycosylation with nigerooligosaccharides as acceptors. Therefore, we synthesized transglycosylation products using Mal_5_-α-*p*NP as a donor, nigerose as an acceptor, and an excess of rAoAgtA in dialyzed culture supernatant and obtained Mal_4_α1,3Glc, Mal_3_α1,3Glc, and Mal_2_α1,3Glc, in which maltooligosaccharide or glucose was transferred to the non-reducing end of nigerose with a formation of an α-1,4-glycosidic bond (Supplementary Tables 2, 3).

### 3-α-Maltooligosylglucose Is a Substrate of Recombinant AoAgtA

The 3-α-maltooligosylglucose motifs are present as part of the speculative cell wall α-1,3-glucan structure. We evaluated the behavior of rAoAgtA with 25 mM Mal_4_α1,3Glc and found that it cleaved Mal_4_α1,3Glc at the first, second, and third glycosidic bonds from the non-reducing end at the 19: 49: 33 ratio (Table 3). The rAoAgtA substrate-degradation velocity for Mal_4_α1,3Glc was 0.133 ± 0.008 mM/min (mean ± standard deviation of three replicates), whereas rAoAgtA scarcely degraded nigeropentaose, which contains only α-1,3-glycosidic bonds. In Mal_5_, the third glycosidic bond from the non-reducing end was mainly cleaved by rAoAgtA, as described in the Analysis of the Modes of Bond Cleavage subsection (Table 3). The rAoAgtA substrate-degradation velocity for Mal_5_ was 0.320 ± 0.027 mM/min, and it was slower for Mal_4_α1,3Glc (0.42-fold, *P* < 0.01 in *t*-test) than for Mal_5_. The rAoAgtA activity for Mal_4_α1,3Glc showed two interesting characteristics: (i) a random cleavage mode of α-1,4-glycosidic bonds for Mal_4_α1,3Glc but not for Mal_5_; (ii) rAoAgtA catalyzed hydrolysis and self-transglycosylation with Mal_5_ but almost exclusively self-transglycosylation with Mal_4_α1,3Glc (data not shown). We found that rAoAgtA randomly cleaves the α-1,4-glycosidic bonds of Mal_4_α1,3Glc. Its structure can be considered as part of the structure of cell wall α-1,3-glucan in *Aspergillus* fungi (Miyazawa et al., 2018).

### The *agtA*-Overexpressing Strains of *Aspergillus nidulans*

Since we have not established a method to measure the MW of the alkali-soluble glucan in the AS2 fractions derived from any *A. oryzae* strains, we introduced the *agtA* overexpression cassette into the Δ*amyD* and *agsB*^*OE*^Δ*amyD* strains of *A. nidulans* to obtain the Δ*amyD-agtA*^*OE*^ and *agsB*^*OE*^Δ*amyD-agtA*^*OE*^ strains (Supplementary Figure 4). In submerged culture, the Δ*amyD* strain formed tightly aggregated hyphal pellets, but the hyphae of the Δ*amyD-agtA*^*OE*^ strain were almost fully dispersed (Figure 4). The latter result is consistent with that for the Δ*amyD-amyD*^*OE*^ strain (Figure 4) reported by Miyazawa et al. (2022). The flask culture of the *agsB*^*OE*^Δ*amyD-agtA*^*OE*^ strain seemed cloudier than those of *agsB*^*OE*^Δ*amyD* and *agsB*^*OE*^Δ*amyD-amyD*^*OE*^ strains, but there was only a slight difference in the hyphal pellets they formed (Figure 4). Overexpression of *agtA* had no marked effect on the phenotype of the *agsB*^*OE*^Δ*amyD* strain.

**Figure 4.**
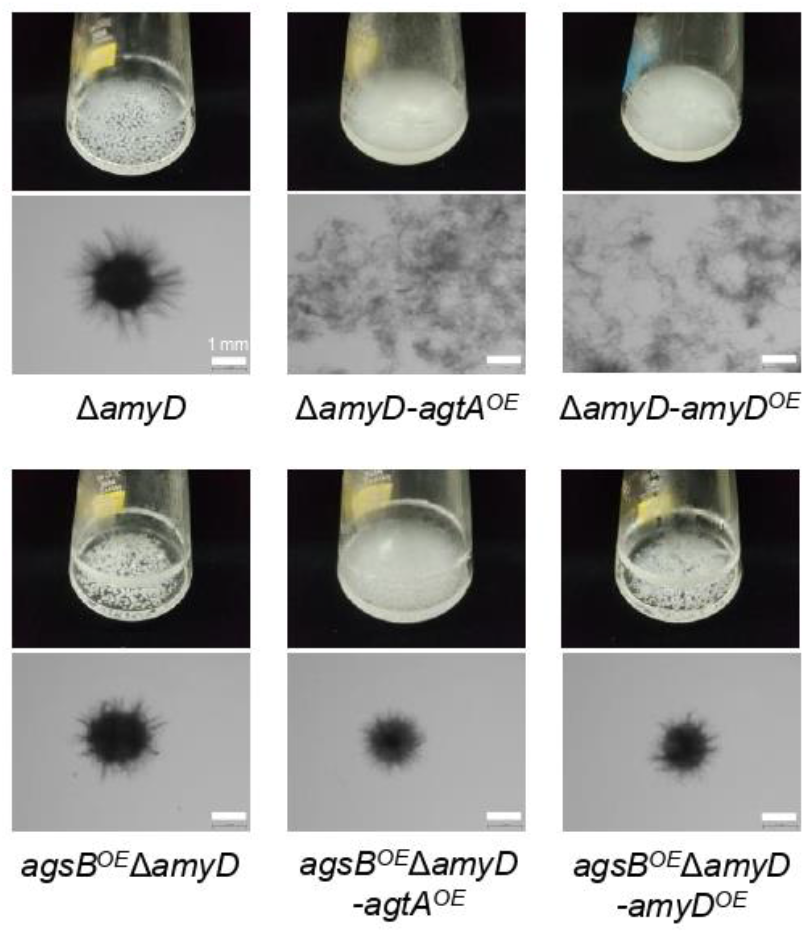
Growth characteristics of *agtA*^*OE*^ strains of *A. nidulans* in submerged culture. Conidia (final concentration, 5 × 10^5^ /mL) of each strain were inoculated into 50 mL of Czapex–Dox (CD) medium and rotated at 160 rpm at 37°C for 24 h. Upper images, cultures in Erlenmeyer flasks; lower images, representative hyphae under a stereomicroscope. Scale bars, 1 mm.

We determined the amount of glucose and MW of glucan in the AS2 fractions from the *agsB*^*OE*^Δ*amyD* and *agsB*^*OE*^Δ*amyD-agtA*^*OE*^ strains. The proportion of glucose in the AS2 fraction seemed to be slightly lower in *agsB*^*OE*^Δ*amyD-agtA*^*OE*^ than in *agsB*^*OE*^Δ*amyD* (Figure 5A), but the difference was not statistically significant, presumably due to the masking effect of *agsB* overexpression. On the other hand, the number-average MW of glucan in the AS2 fraction from *agsB*^*OE*^Δ*amyD-agtA*^*OE*^ (95,700 ± 1,300) was significantly lower than that from *agsB*^*OE*^Δ*amyD* (462,000 ± 38,000; Figure 5B; Table 5, *P* < 0.01 in Welch’s test). These results indicate that AoAgtA decreases the MW of cell wall α-1,3-glucan, similar to *A. nidulans* AmyD (Miyazawa et al., 2022).

**Figure 5.**
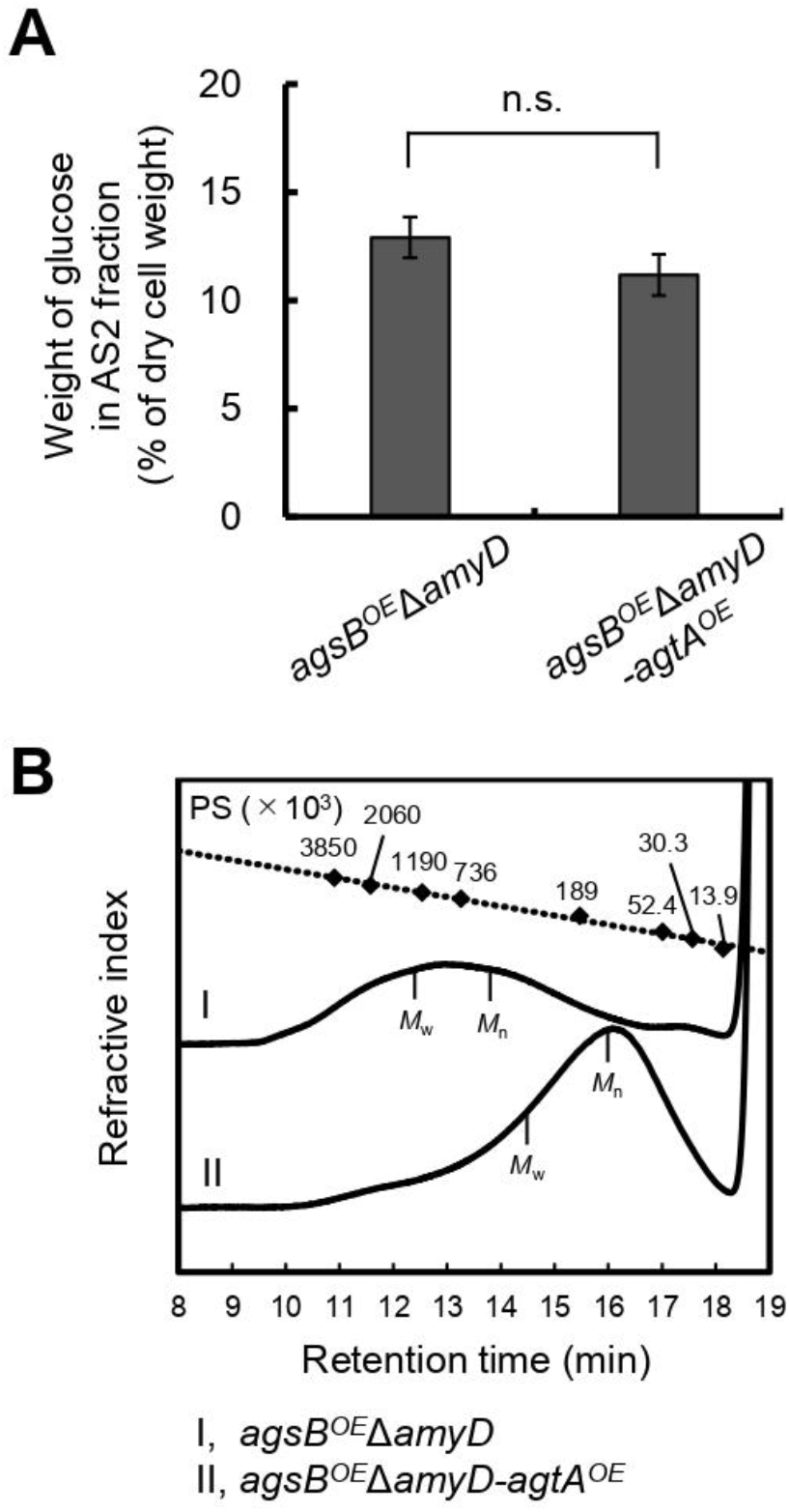
(A) Glucose content and (B) gel permeation chromatography elution profile of the AS2 fraction in the *agsB*^*OE*^Δ*amyD-agtA*^*OE*^ strain of *A. nidulans*. **(A)** Conidia (5 × 10^5^ /mL) of each strain were inoculated into CD medium and rotated at 160 rpm at 37°C for 24 h. Error bars represent the standard error of the mean calculated from three replicates. n.s., not significant in *t*-test (*P* ≥ 0.05). **(B)** The AS2 fraction from 24-h-cultured mycelia of each strain was dissolved in 20 mM LiCl/*N,N*-dimethylacetamide. The elution profile was monitored with a refractive index detector. Molecular weights (MW) of the glucan peaks were determined from a calibration curve of polystyrene (PS) standards (♦). *M*_w_, weight-average MW; *M*_n_, number-average MW.

**Table 5.** Molecular weight of alkali-soluble glucan from the cell wall.

| Sample | $M_p^b$ | $M_w^c$ | $M_n^d$ | $M_w/M_n$ |
| --- | --- | --- | --- | --- |
| *agsB<sup>OE</sup>ΔamyD* AS2<sup>a</sup> | 771 000 ± 106 000 | 1 330 000 ± 80 000 | 462 000 ± 38 000 | 2.88 ± 0.07 |
| *agsB<sup>OE</sup>ΔamyD-agtA<sup>OE</sup>* AS2 | 90 700 ± 4 300 | 285 000 ± 14 000 | 95 700 ± 1 300 | 2.97 ± 0.11 |

<sup>a</sup>AS2, insoluble components after dialysis of the alkali-soluble fraction.
<sup>b</sup> Peak molecular weight.
<sup>c</sup> Weight-average molecular weight.
<sup>d</sup> Number-average molecular weight.
data are mean ± standard deviation of three replicates.

## DISCUSSION

Although Agt proteins encoded by the orthologous *agtA* genes in *Aspergillus* fungi are thought to be GPI-anchored α-amylases, their function may be related to the biosynthesis of cell wall α-1,3-glucan rather than to starch catabolism (van der Kaaij et al., 2007; He et al., 2014, 2017; Miyazawa et al., 2022). In the present study, similarly to *amyD* in *A. nidulans* (He et al., 2014, 2017; Miyazawa et al., 2022), *agtA* overexpression in *A. oryzae* decreased the amount of α-1,3-glucan in the cell wall (Figure 2), suggesting that AoAgtA suppresses cell wall α-1,3-glucan biosynthesis in *A. oryzae*. We also investigated the enzymatic properties of rAoAgtA to further understand its contribution to cell wall α-1,3-glucan biosynthesis.

Among several types of glucans, rAoAgtA degraded only soluble starch (Supplementary Table 6). Therefore, AoAgtA appeared to act on α-1,4-glycosidic linkages and may degrade the spacer composed of α-1,4-linked glucose residues that is thought to be incorporated into cell wall α-1,3-glucan (Figure 1) during its biosynthesis. We analyzed the modes of bond cleavage by rAoAgtA, including hydrolysis and transglycosylation, using maltooligosaccharide substrates. rAoAgtA showed an endo-type cleavage mode for substrates with a DP at least 5 (Table 3). The weak activity of rAoAgtA on soluble starch and its substrate specificity for maltooligosaccharide substrates were consistent with the properties of AnAgtA of *A. niger*, which also has 4-α-glucanotransferase activity and is expected to belong to a new subgroup of GH13 (van der Kaaij et al., 2007). Although the length (DP) of the spacer has not been clarified, if AoAgtA cleaves the spacer, its DP should be at least 5, as suggested by the failure of rAoAgtA to degrade nigeran, a polysaccharide consisting of alternating α-1,3- and α-1,4-linked glucose residues (Supplementary Table 6), and by the modes of bond cleavage in maltooligosaccharide substrates (Table 3). However, the kinetic studies of PNM degradation (Table 4) demonstrated an extremely low rAoAgtA activity for maltooligosaccharide substrates compared with other known α-amylases. The reason for such low activity may be a replacement of a His residue in Region I of the Agt proteins, which was also mentioned in the case of AnAgtA by van der Kaaij et al. (2007) (Supplementary Table 7). This residue is particularly conserved among the members of the α-amylase family (Svensson, 1994; Kuriki et al., 2006), and its mutation reportedly decreases enzyme activity and alters the type of reaction catalyzed by the enzyme (Nakamura et al., 1993; Svensson, 1994; Chang et al., 2003; Leemhuis et al., 2004). This His residue is changed to Asn in AoAgtA (Supplementary Table 7). The enzymatic characteristics of rAoAgtA might reflect the evolutionary divergence of the functions of Agt proteins from those of α-amylases highly active on starch.

We consider that Agt proteins are specialized in cleaving α-1,4-linked oligosaccharides or glucan with α-1,3-glycosidic bonds on the side of the reducing end or both sides *in vivo*. As preparation of such predicted substrates is currently difficult, we examined the effect of nigerooligosaccharides on rAoAgtA catalytic activity and found that they increased Mal_5_-α-*p*NP-degradation (Supplementary Figure 9). AnAgtA can use nigerooligosaccharides with a small DP as acceptors for transglycosylation (van der Kaaij et al., 2007). We predicted that rAoAgtA also conducts transglycosylation with nigerooligosaccharides as acceptors. In fact, 3-α-maltooligosylglucose was synthesized by rAoAgtA transglycosylation activity with Mal_5_-α-*p*NP as a donor and nigerose as an acceptor (Supplementary Tables 2, 3). The transfer of maltooligosaccharide to nigerooligosaccharide with a formation of an α-1,4-glycosidic bond, which is catalyzed by rAoAgtA, has not been known for other α-amylases. The study of AnAgtA has not elucidated the chemical structures of the transglycosylation products with nigerooligosaccharides as acceptors (van der Kaaij et al., 2007). The degradation of Mal_5_-α-*p*NP catalyzed by rAoAgtA likely proceeds *via* a covalent glycosyl (mainly Mal_3_)–enzyme intermediate. We expect that such an intermediate would be more vulnerable to nucleophilic attack with oligosaccharides than with H_2_O.

The structure of 3-α-maltooligosylglucose containing both nigerooligosaccharide and maltooligosaccharide components is present as parts of the structure in the speculative cell wall α-1,3-glucan structure. In fact, rAoAgtA was able to cleave Mal_4_α1,3Glc (Table 3). The activity of rAoAgtA with Mal_4_α1,3Glc was weak, with a slower substrate-degradation velocity than that for Mal_5_. The length of the maltooligosaccharide moiety of the Mal_4_α1,3Glc might be insufficient for the maximum activity of rAoAgtA, as mentioned above. Interestingly, we found that the rAoAgtA mode of bond cleavage in Mal_4_α1,3Glc clearly differed from that in Mal_5_ (Table 3).

We here propose a hypothetical role of AoAgtA in cell wall α-1,3-glucan biosynthesis. rAoAgtA randomly cleaved only the α-1,4-glycosidic bonds of Mal_4_α1,3Glc (Table 3), and the structure of Mal_4_α1,3Glc is similar to the predicted boundary structure between α-1,3-glucan main chain and the spacer in cell wall α-1,3-glucan synthesized by AgsB (Figure 6A). Miyazawa et al. (2022) suggested that AmyD of *A. nidulans* requires a GPI anchor to act on α-1,3-glucan *in vivo* and that AmyD reacts with cell wall α-1,3-glucan shortly after it is synthesized by α-1,3-glucan synthase on the plasma membrane. Therefore, we speculate that AoAgtA cleaves some of the spacers of cell wall α-1,3-glucan in the process of synthesis *in vivo* (Figure 6B). The random mode of α-1,4-glycosidic bond cleavage in Mal_4_α1,3Glc by rAoAgtA (Table 3) supports the possibility that AoAgtA cleaves spacers in cell wall α-1,3-glucan. In fact, the MW of glucan in the AS2 fraction derived from the *agtA*^*OE*^ strain of *A. nidulans* was significantly lower than in that from the control stain (Figure 5B; Table 5). Thus, similar to AmyD (Miyazawa et al., 2022), AoAgtA decreased the MW of cell wall α-1,3-glucan *in vivo*. The MW of glucan in the AS2 fraction extracted from *agsB*^*OE*^Δ*amyD* was not affected by purified rAoAgtA (data not shown), perhaps because mature cell wall α-1,3-glucan is packed by hydrogen bonds, and its spacers cannot be accessed by the enzyme. To prove that AoAgtA acts on cell wall α-1,3-glucan *in vitro*, it is necessary to establish a new evaluation system.

**Figure 6.**
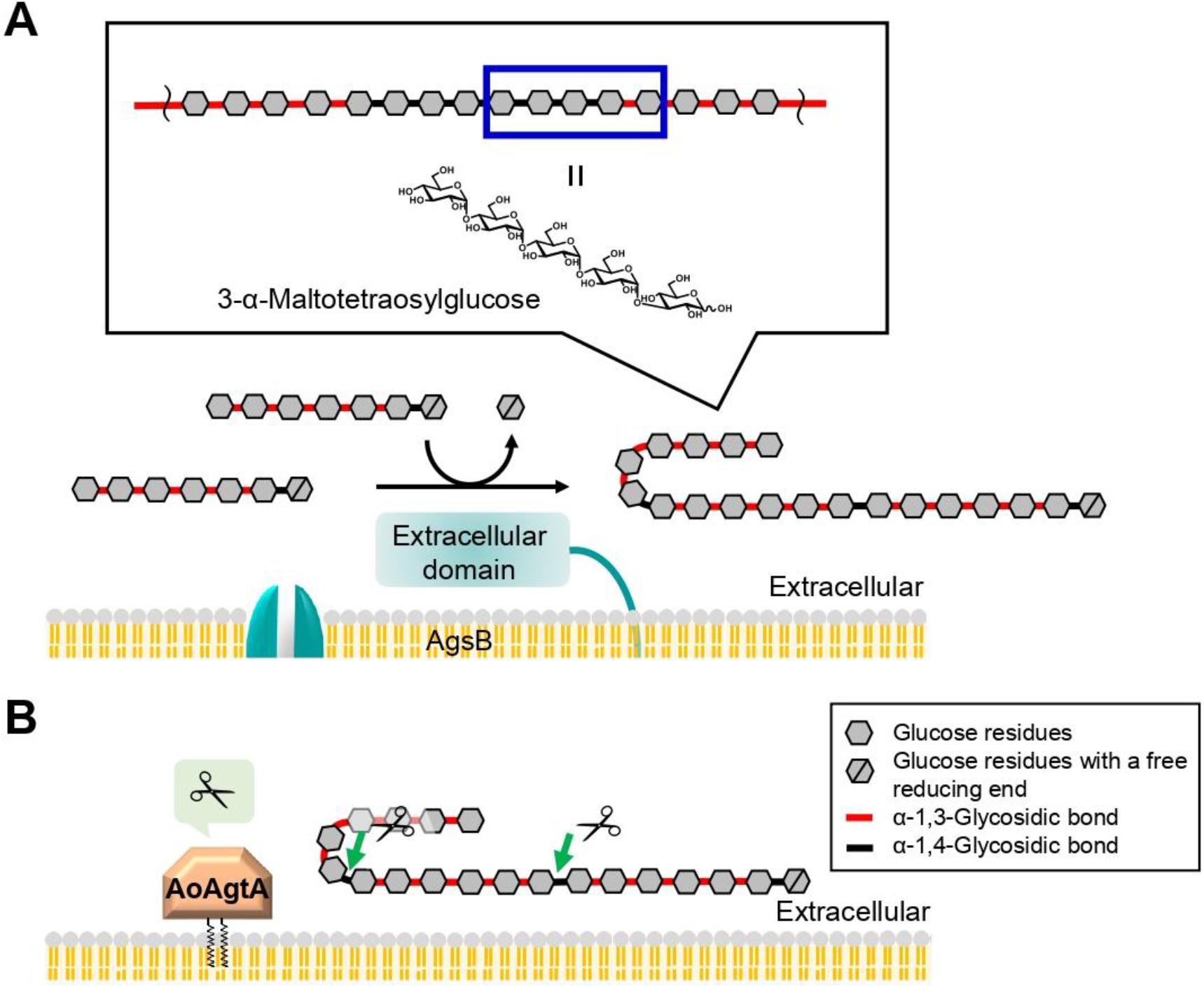
Predicted role of AoAgtA in cell wall α-1,3-glucan biosynthesis in *A. oryzae*. **(A)** During biosynthesis, α-1,3-glucan units may be interconnected by transglycosylation catalyzed by the extracellular domain of AgsB; the structure of 3-α-maltotetraosylglucose is consistent with the structure in which α-1,3-glucan units are linked. **(B)** We speculate that AoAgtA cleaves the spacers composed of α-1,4-linked glucose residues in cell wall α-1,3-glucan before its insolubilization.

In the present study, we showed that overexpression of the *agtA* gene in *A. oryzae* decreased the amount of α-1,3-glucan in the cell wall, suggesting that AoAgtA suppresses cell wall α-1,3-glucan biosynthesis. Analysis using the *agtA*^*OE*^ strain of *A. nidulans* showed that AoAgtA decreases the MW of cell wall α-1,3-glucan. Purified rAoAgtA randomly cleaved the α-1,4-glycosidic bonds of Mal_4_α1,3Glc, which can be considered as part of the structure of cell wall α-1,3-glucan. We conclude that AoAgtA likely cleaves the spacers composed of α-1,4-linked glucose residues in cell wall α-1,3-glucan before its insolubilization. Taken together with the results of *in vivo* functional analysis of the *amyD* gene of *A. nidulans* (Miyazawa et al., 2022), the present study suggests that AoAgtA plays an important role in the biosynthesis of cell wall α-1,3-glucan.

## DATA AVAILABILITY STATEMENT

The original contributions presented in the study are included in the article/Supplementary Material, further inquiries can be directed to the corresponding author.

## AUTHOR CONTRIBUTIONS

AK, KM, AY, and KA conceived and designed the experiments. AK performed most experiments and analyzed the data. AK and MS constructed the *A. oryzae* mutants. AK, YT, SY, and MS constructed the *P. pastoris* strain and purified rAoAgtA. AK and MO performed enzymatic assays and chemical analyses of oligosaccharides. AK and KM constructed the *A. nidulans* mutants. TN and HN produced nigerooligosaccharides. SK and TI enzymatically produced bacterial α-1,3-glucan. MH performed structural analysis of AoAgtA. KA supervised this research and acquired funding. AK, KM, MO, AY, and KA wrote the paper. All authors contributed to the article and approved the submitted version.

## ACKNOWLEDGMENTS

We are grateful to Kikkoman for providing maltooligosaccharides. We thank Professor Emeritus Tasuku Nakajima (Tohoku University) for providing pullulan and for critical discussion on the research, Dr. Keiko Uechi (University of the Ryukyus) for providing nigeran, and Dr. Daisuke Sugimori (Fukushima University) for lending equipment.

## REFERENCES

Beauvais, A., Bozza, S., Kniemeyer, O., Formosa, C., Balloy, V., Henry, C., et al. (2013). Deletion of the α-(1,3)-glucan synthase genes induces a restructuring of the conidial cell wall responsible for the avirulence of Aspergillus fumigatus. PLoS Pathog. 9:e1003716. doi: 10.1371/journal.ppat.1003716.

Chang, C. T., Lo, H. F., Chi, M. C., Yao, C. Y., Hsu, W. H., and Lin, L. L. (2003). Identification of essential histidine residues in a recombinant α-amylase of thermophilic and alkaliphilic Bacillus sp. strain TS-23. Extremophiles 7, 505–509. doi: 10.1007/s00792-003-0341-8.

Choma, A., Wiater, A., Komaniecka, I., Paduch, R., Pleszczyńska, M., and Szczodrak, J. (2013). Chemical characterization of a water insoluble (1 → 3)-α-D-glucan from an alkaline extract of Aspergillus wentii. Carbohydr. Polym. 91, 603–608. doi: 10.1016/j.carbpol.2012.08.060.

Czerwonka, A., Wiater, A., Komaniecka, I., Adamczyk, P., Rzeski, W., and Pleszczyńska, M. (2019). Antitumour effect of glucooligosaccharides obtained via hydrolysis of α-(1 → 3)-glucan from Fomitopsis betulina. Mol. Biol. Rep. 46, 5977–5982. doi: 10.1007/s11033-019-05032-x.

Doner, L. W., and Irwin, P. L. (1992). Assay of reducing end-groups in oligosaccharide homologues with 2,2’-bicinchoninate. Anal. Biochem. 202, 50–53. doi: 10.1016/0003-2697(92)90204-k.

Fujikawa, T., Kuga, Y., Yano, S., Yoshimi, A., Tachiki, T., Abe, K., et al. (2009). Dynamics of cell wall components of Magnaporthe grisea during infectious structure development. Mol. Microbiol. 73, 553–570. doi: 10.1111/j.1365-2958.2009.06786.x.

Fujikawa, T., Sakaguchi, A., Nishizawa, Y., Kouzai, Y., Minami, E., Yano, S., et al. (2012). Surface α-1,3-glucan facilitates fungal stealth infection by interfering with innate immunity in plants. PLoS Pathog. 8:e1002882. doi: 10.1371/journal.ppat.1002882.

Grün, C. H., Hochstenbach, F., Humbel, B. M., Verkleij, A. J., Sietsma, J. H., Klis, F. M., et al. (2005). The structure of cell wall α-glucan from fission yeast. Glycobiology 15, 245–257. doi: 10.1093/glycob/cwi002.

Hara, S., Tsuji, R. F., Hatamoto, O., and Masuda, T. (2002). A simple method for enrichment of uninucleate conidia of Aspergillus oryzae. Biosci. Biotechnol. Biochem. 66, 693–696. doi: 10.1271/bbb.66.693.

Hattori, T., Kato, Y., Uno, S., and Usui, T. (2013). Mode of action of a β-(1→6)-glucanase from Penicillium multicolor. Carbohydr. Res. 366, 6–16. doi: 10.1016/j.carres.2012.11.002.

He, X., Li, S., and Kaminskyj, S. (2017). An amylase-like protein, AmyD, is the major negative regulator for α-glucan synthesis in Aspergillus nidulans during the asexual life cycle. Int. J. Mol. Sci. 18:695. doi: 10.3390/ijms18040695.

He, X., Li, S., and Kaminskyj, S. G. W. (2014). Characterization of Aspergillus nidulans α-glucan synthesis: roles for two synthases and two amylases. Mol. Microbiol. 91, 579–595. doi: 10.1111/mmi.12480.

Ichikawa, H., Miyazawa, K., Komeiji, K., Susukida, S., Zhang, S., Muto, K., et al. (2022). Improved recombinant protein production in Aspergillus oryzae lacking both α-1,3-glucan and galactosaminogalactan in batch culture with a lab-scale bioreactor. J. Biosci. Bioeng. 133, 39– 45. doi: 10.1016/j.jbiosc.2021.09.010.

Jumper, J., Evans, R., Pritzel, A., Green, T., Figurnov, M., Ronneberger, O., et al. (2021). Highly accurate protein structure prediction with AlphaFold. Nature 596, 583–589. doi: 10.1038/s41586-021-03819-2.

Kazim, A. R. S., Jiang, Y., Li, S., and He, X. (2021). Aspergillus nidulans AmyG functions as an intracellular α-amylase to promote α-glucan synthesis. Microbiol. Spectr. 9:e00644–21. doi: 10.1128/Spectrum.00644-21.

Kuriki, T., Takata, H., Yanase, M., Ohdan, K., Fujii, K., Terada, Y., et al. (2006). The concept of the α-amylase family: a rational tool for interconverting glucanohydrolases/glucanotransferases, and their specificities. J. Appl. Glycosci. 53, 155–161. doi: 10.5458/jag.53.155.

Laemmli, U. K. (1970). Cleavage of structural proteins during the assembly of the head of bacteriophage T4. Nature 227, 680–685. doi: 10.1038/227680a0.

Latgé, J. P. (2010). Tasting the fungal cell wall. Cell. Microbiol. 12, 863–872. doi: 10.1111/j.1462-5822.2010.01474.x.

Leemhuis, H., Wehmeier, U. F., and Dijkhuizen, L. (2004). Single amino acid mutations interchange the reaction specificities of cyclodextrin glycosyltransferase and the acarbose-modifying enzyme acarviosyl transferase. Biochemistry 43, 13204–13213. doi: 10.1021/bi049015q.

Minetoki, T., Tsuboi, H., Koda, A., and Ozeki, K. (2003). Development of high expression system with the improved promoter using the cis-acting element in Aspergillus species. J. Biol. Macromol. 3, 89–96.

Miyazawa, K., Yamashita, T., Takeuchi, A., Kamachi, Y., Yoshimi, A., Tashiro, Y., et al. (2022). A glycosylphosphatidylinositol-anchored α-amylase encoded by amyD contributes to a decrease in the molecular mass of cell wall α-1,3-glucan in Aspergillus nidulans. Front. Fungal Biol. 2:821946. doi: 10.3389/ffunb.2021.821946.

Miyazawa, K., Yoshimi, A., and Abe, K. (2020). The mechanisms of hyphal pellet formation mediated by polysaccharides, α-1,3-glucan and galactosaminogalactan, in Aspergillus species. Fungal Biol. Biotechnol. 7:10. doi: 10.1186/s40694-020-00101-4.

Miyazawa, K., Yoshimi, A., Kasahara, S., Sugahara, A., Koizumi, A., Yano, S., et al. (2018). Molecular mass and localization of α-1,3-glucan in cell wall control the degree of hyphal aggregation in liquid culture of Aspergillus nidulans. Front. Microbiol. 9:2623. doi: 10.3389/fmicb.2018.02623.

Miyazawa, K., Yoshimi, A., Sano, M., Tabata, F., Sugahara, A., Kasahara, S., et al. (2019). Both galactosaminogalactan and α-1,3-glucan contribute to aggregation of Aspergillus oryzae hyphae in liquid culture. Front. Microbiol. 10:2090. doi: 10.3389/fmicb.2019.02090.

Miyazawa, K., Yoshimi, A., Zhang, S., Sano, M., Nakayama, M., Gomi, K., et al. (2016). Increased enzyme production under liquid culture conditions in the industrial fungus Aspergillus oryzae by disruption of the genes encoding cell wall α-1,3-glucan synthase. Biosci. Biotechnol. Biochem. 80, 1853–1863. doi: 10.1080/09168451.2016.1209968.

Mizutani, O., Kudo, Y., Saito, A., Matsuura, T., Inoue, H., Abe, K., et al. (2008). A defect of LigD (human Lig4 homolog) for nonhomologous end joining significantly improves efficiency of gene-targeting in Aspergillus oryzae. Fungal Genet. Biol. 45, 878–889. doi: 10.1016/j.fgb.2007.12.010.

Nakamura, A., Haga, K., and Yamane, K. (1993). Three histidine residues in the active center of cyclodextrin glucanotransferase from alkalophilic Bacillus sp. 1011: effects of the replacement on pH dependence and transition-state stabilization. Biochemistry 32, 6624–6631. doi: 10.1021/bi00077a015.

Nihira, T., Nishimoto, M., Nakai, H., Ohtsubo, K., and Kitaoka, M. (2014). Characterization of two α-1,3-glucoside phosphorylases from Clostridium phytofermentans. J. Appl. Glycosci. 61, 59– 66. doi: 10.5458/jag.jag.JAG-2013_013.

Nitta, Y., Mizushima, M., Hiromi, K., and Ono, S. (1971). Influence of molecular structures of substrates and analogues on Taka-amylase A catalyzed hydrolyses. I. Effect of chain length of linear substrates. J. Biochem. 69, 567–576.

Okada, M., Ogawa, K., and Usui, T. (2000). Substrate specificity of human and bovine pancreatic α-amylases using end-blocked maltooligosaccharides. J. Appl. Glycosci. 47, 35–44. doi: 10.5458/jag.47.35.

Puanglek, S., Kimura, S., Enomoto-Rogers, Y., Kabe, T., Yoshida, M., Wada, M., et al. (2016). In vitro synthesis of linear α-1,3-glucan and chemical modification to ester derivatives exhibiting outstanding thermal properties. Sci. Rep. 6:30479. doi: 10.1038/srep30479.

Rappleye, C. A., Eissenberg, L. G., and Goldman, W. E. (2007). Histoplasma capsulatum α-(1,3)-glucan blocks innate immune recognition by the β-glucan receptor. Proc. Natl. Acad. Sci. U. S. A. 104, 1366–1370. doi: 10.1073/pnas.0609848104.

Rappleye, C. A., Engle, J. T., and Goldman, W. E. (2004). RNA interference in Histoplasma capsulatum demonstrates a role for α-(1,3)-glucan in virulence. Mol. Microbiol. 53, 153–165. doi: 10.1111/j.1365-2958.2004.04131.x.

Suganuma, T. (1983). Subsite structures and transglycosylation of Taka-amylase A and Bacillus subtilis saccharifying α-amylase. J. Jpn. Soc. Starch Sci. 30, 149–158.

Svensson, B. (1994). Protein engineering in the α-amylase family: catalytic mechanism, substrate specificity, and stability. Plant Mol. Biol. 25, 141–157. doi: 10.1007/BF00023233.

Tamano, K., Satoh, Y., Ishii, T., Terabayashi, Y., Ohtaki, S., Sano, M., et al. (2007). The β-1,3-exoglucanase gene exgA (exg1) of Aspergillus oryzae is required to catabolize extracellular glucan, and is induced in growth on a solid surface. Biosci. Biotechnol. Biochem. 71, 926–934. doi: 10.1271/bbb.60591.

Tunyasuvunakool, K., Adler, J., Wu, Z., Green, T., Zielinski, M., žídek, A., et al. (2021). Highly accurate protein structure prediction for the human proteome. Nature 596, 590–596. doi: 10.1038/s41586-021-03828-1.

Usui, T., and Murata, T. (1988). Enzymatic synthesis of p-nitrophenyl α-maltopentaoside in an aqueous-methanol solvent system by maltotetraose-forming amylase: a substrate for human amylase in serum. J. Biochem. 103, 969–972. doi: 10.1093/oxfordjournals.jbchem.a122395.

Usui, T., Ogawa, K., Nagai, H., and Matsui, H. (1992). Enzymatic synthesis of p-nitrophenyl 45-O-β-D-galactosyl-α-maltopentaoside as a substrate for human α-amylases. Anal. Biochem. 202, 61– 67. doi: 10.1016/0003-2697(92)90206-m.

Utsumi, Y., Yoshida, M., Francisco, P. B. J., Sawada, T., Kitamura, S., and Nakamura, Y. (2009). Quantitative assay method for starch branching enzyme with bicinchoninic acid by measuring the reducing terminals of glucans. J. Appl. Glycosci. 56, 215–222. doi: 10.5458/jag.56.215.

van der Kaaij, R. M., Yuan, X. L., Franken, A., Ram, A. F. J., Punt, P. J., van der Maarel, M. J. E. C., et al. (2007). Two novel, putatively cell wall-associated and glycosylphosphatidylinositol-anchored α-glucanotransferase enzymes of Aspergillus niger. Eukaryot. Cell 6, 1178–1188. doi: 10.1128/EC.00354-06.

Vujičić-žagar, A., and Dijkstra, B. W. (2006). Monoclinic crystal form of Aspergillus niger α-amylase in complex with maltose at 1.8 Å resolution. Acta Crystallogr. Sect. F: Struct. Biol. Cryst. Commun. 62, 716–721. doi: 10.1107/S1744309106024729.

Yoshimi, A., Miyazawa, K., and Abe, K. (2016). Cell wall structure and biogenesis in Aspergillus species. Biosci. Biotechnol. Biochem. 80, 1700–1711. doi: 10.1080/09168451.2016.1177446.

Yoshimi, A., Miyazawa, K., and Abe, K. (2017). Function and biosynthesis of cell wall α-1,3-glucan in fungi. J. Fungi 3:63. doi: 10.3390/jof3040063.

Yoshimi, A., Sano, M., Inaba, A., Kokubun, Y., Fujioka, T., Mizutani, O., et al. (2013). Functional analysis of the α-1,3-glucan synthase genes agsA and agsB in Aspergillus nidulans: AgsB is the major α-1,3-glucan synthase in this fungus. PLoS ONE 8:e54893. doi: 10.1371/journal.pone.0054893.

Zhang, S., Sato, H., Ichinose, S., Tanaka, M., Miyazawa, K., Yoshimi, A., et al. (2017). Cell wall α-1,3-glucan prevents α-amylase adsorption onto fungal cell in submerged culture of Aspergillus oryzae. J. Biosci. Bioeng. 124, 47–53. doi: 10.1016/j.jbiosc.2017.02.013.

