## Supplementary Material for "Cleavage of α-1,4-Glycosidic Linkages by the Glycosylphosphatidylinositol-Anchored α-Amylase AgtA Decreases the Molecular Weight of Cell Wall α-1,3-Glucan in *Aspergillus oryzae*"

### MATERIALS AND METHODS

#### Enzymatic Synthesis of *p*-Nitrophenyl α-Maltooligosides

A mixture (530 μL) containing $Mal_5$-α-*p*NP (100 mg, 200 mM) and 180 mU/mL rAoAgtA in dialyzed culture supernatant in 50 mM Na-Ac buffer (pH 5.5) was incubated at 40°C for 24 h. The reaction was stopped by adding 11 mL of methanol. The reaction mixture was dried, dissolved in a small amount of $H_2O$, and then applied to a Toyopearl HW-40S column (4.0 × 60 cm) equilibrated with $H_2O$ at a flow rate of 0.6 mL/min. The eluate was collected in 10-mL fractions (950 mL in total). Each fraction was analyzed by HPLC. The HPLC system consisted of a Mightysil Si60 column (4.6 × 250 mm; Kanto Chemical, Chuo, Japan) and a Jasco Intelligent System Liquid Chromatograph; detection was performed at 300 nm. The bound material was eluted with 75% acetonitrile at a flow rate of 1.0 mL/min at 40°C. Fractions corresponding to $Mal_{2–8}$-α-*p*NP were concentrated and then lyophilized. Products with a purity of less than 98% were rechromatographed under the same conditions. As a result, $Mal_8$-α-*p*NP (4.7 mg, yield 3.1%), $Mal_7$-α-*p*NP (5.5 mg, 4.1%), $Mal_6$-α-*p*NP (7.1 mg, 6.1%), $Mal_5$-α-*p*NP (12.3 mg, 12.3%), $Mal_4$-α-*p*NP (6.2 mg, 7.5%), $Mal_3$-α-*p*NP (10.8 mg, 16.4%), and $Mal_2$-α-*p*NP (11.9 mg, 24.4%) were obtained. The structures of the synthesized $Mal_{2–8}$-α-*p*NP were evaluated by $^1$H NMR analysis in $D_2O$ (Supplementary Figure 3); 500 MHz $^1$H NMR spectra were recorded using a Jeol ECX-500 II spectrometer (Jeol, Akishima, Japan).

#### Biochemical Characterization of Recombinant AoAgtA

Purified rAoAgtA was used in all tests except that for pH stability, where dialyzed culture supernatant was used.

The optimum temperature was determined within a range of 4–60°C in 50 mM Na-Ac buffer (pH 5.5) (Supplementary Figure 6A). To measure thermostability, the enzyme solutions were heated at 10–70°C for 30 min and then residual rAoAgtA activity was measured (Supplementary Figure 6B).

The optimum pH was determined at 40°C in the following 50 mM buffers: glycine-HCl (pH 1–3), citric acid-NaOH (pH 3–4), Na-Ac (pH 4–6), 3-morpholinopropanesulfonic acid (MOPS)-NaOH (pH 6–7), Tris-HCl (pH 7–9), and glycine-NaOH (pH 9–10) (Supplementary Figure 6C). To measure pH stability, enzyme solutions were incubated at 4°C for 30 min in 10 mM each buffer mentioned above and then residual rAoAgtA activity was measured (Supplementary Figure 6D).

The effect of metal ions (LiCl, KCl, $MgCl_2$, $CaCl_2$, $MnCl_2$, $FeCl_3$, $CoCl_2$, $CuCl_2$, $ZnCl_2$, and $AlCl_3$) and EDTA on rAoAgtA activity was evaluated by measuring rAoAgtA activity in 50 mM Na-Ac buffer (pH 5.5) in the presence of 2.5 mM of each metal ion or EDTA; rAoAgtA activity in their absence was considered 100% (Supplementary Table 4).

We also evaluated the effect of $CaCl_2$ on rAoAgtA activity after treatment with EDTA (Supplementary Table 5). The enzyme (0.19 mU) and 25 mM EDTA (2 μL; 2.5 mM at final concentration) were incubated in 63 mM Na-Ac buffer (pH 5.5; 50 mM at final concentration) at 4°C for 30 min without the substrate. Next, the substrate ($Mal_5$-α-*p*NP) was added, and rAoAgtA activity was measured. Alternatively, 25 mM $CaCl_2$ (2 μL; 2.5 mM at final concentration) was added after the incubation with EDTA, and the sample was incubated at 4°C for another 30 min. $Mal_5$-α-*p*NP was then added, and rAoAgtA activity was measured.

**Supplementary Table 1. Primers used in this study.**

| Purpose | Primer name | Sequence (5′ to 3′) |
|---|---|---|
| Construction of *agtA*$^{OE}$ and Δ*agtA* strains of *Aspergillus oryzae* | | |
| | agtA-Fw-NotI | GACAAGCTTGCGGCCGCATGGTTTCGTCGTCATCCCT |
| | agtA-Rv-NotI | AGTCACGTGGCGGCCGCCTACAACAATACCGCAACAAGAC |
| | agtA-up-Fw | TCCTTCCAACACCGATCCAG |
| | agtA-mid-Rv | TGCTGGCGTCGGTACATACC |
| | agtA-LU | CCTTCTTTTCCCGTCCTT |
| | agtA-LL+adeA | ATATACCGTGACTTTTTAGTGAAAGTAACTGGAGTCGT |
| | agtA-RU+adeA | AGTTTCGTCGAGATACTGCCGGAGAAGTTCTTCAGCA |
| | agtA-RL | TCGGCTAAGTTACAGACAG |
| | agtA-AU | GACTCCAGTTACTTTCACTAAAAAGTCACGGTATATCATGAC |
| | agtA-AL | TGCTGAAGAACTTCTCCGGCAGTATCTCGACGAAACTACCTAA |
| Construction of the rAoAgtA expression system | | |
| | agtA-Fw-NdeI | GGAATTCCATATGGTTTCGTCGTCATCCCT |
| | agtA-Rv-SmaI | CCCCCGGGCGCCGCGCCCTTGGTCTTCAG |
| | agtA-Fw-PstI | ATCAGCCGCTGCAGCAACCACAGCAGAATGGAAG |
| | agtA-Rv-XbaI | CCAGTGTGTCTAGATTCGCCCTTTTATTAATGA |
| Construction of heterologous *agtA*$^{OE}$ strains of *Aspergillus nidulans* | | |
| | agtA-dIntron-Fw | TATGGCGCTTGCTAAGAATGTTTTAACCTTCAC |
| | agtA-dIntron-Rv | TTAGCAAGCGCCATATCATCAGTCATACTAGCG |
| | agtA-IF-top-Fw | CGCACCACCTTCAAAATGGTTTCGTCGTCATCCCTG |
| | TagdA-IF-tail-Rv | TTGTGCTTCTCTGCAAGGTGTACGCTTGGTAAAGTTG |
| | AopyrG-IF-Right-Fw | TGCAGAGAAGCACAATTTCCTCATC |
| | Ptef1-tail-Rv | TTTGAAGGTGGTGCGAACTTTGTAG |
| | 397-5 | GAGGCCACTCAGGCCGATATCACC |
| | ANamyD-up-Fw | AGGTTCAACGATCGAACCCAGCAAC |
| | Hph-top-Rv | CCAGCTTGTGTTCCCGGTCTG |
| | AopyrG-IF-Right-Rv | GCCAGTGAATTCGAGCTCAACTGCACCTCAGAAGAAAAGGATG |

**Supplementary Table 2.** **$^{1}$H-chemical shifts and coupling constants of the anomeric protons of 3-α-maltosyl-, 3-α-maltotriosyl-, and 3-α-maltotetraosyl-glucose dissolved in $D_2O$ (25°C).**

| Compound | H1 | | H1' | | H1'' | | H1''' | | H1'''' | |
|---|---|---|---|---|---|---|---|---|---|---|
| | **α** | **β** | **α** | **β** | **α** | **β** | **α** | **β** | **α** | **β** |
| | ***J*, Hz** | ***J*, Hz** | ***J*, Hz** | ***J*, Hz** | ***J*, Hz** | ***J*, Hz** | ***J*, Hz** | ***J*, Hz** | ***J*, Hz** | ***J*, Hz** |
| 3-α-Maltosylglucose (Koto et al., 1992) | 5.15 | 4.59 | 5.28 | 5.30 | 5.33 | 5.33 | | | | |
| | 3.5 | 8.0 | 4.0 | 4.0 | 3.5 | 3.5 | | | | |
| 3-α-Maltosylglucose | 5.16 | 4.59 | 5.29 | 5.30 | 5.34 | 5.34 | | | | |
| | 3.8 | 8.1 | 3.9 | 3.9 | 3.9 | 3.9 | | | | |
| 3-α-Maltotriosylglucose | 5.16 | 4.59 | 5.29 | 5.31 | 5.33 | 5.33 | 5.32 | 5.32 | | |
| | 3.8 | 8.0 | 3.9 | 3.9 | 4.0 | 4.0 | 4.0 | 4.0 | | |
| 3-α-Maltotetraosylglucose | 5.16 | 4.59 | 5.29 | 5.31 | 5.34 | 5.34 | 5.32 | 5.32 | 5.31 | 5.31 |
| | 3.8 | 8.0 | 3.9 | 3.9 | 4.0 | 4.0 | 3.9 | 3.9 | 3.9 | 3.9 |

For each compound: upper row, $^{1}$H-chemical shifts; lower row, coupling constants (Hz).

**Supplementary Table 3. $^{13}$C-chemical shifts of 3-α-maltosyl-, 3-α-maltotriosyl-, and 3-α-maltotetraosyl-glucose dissolved in $D_2O$ (25°C).**

| Carbon number | Compounds | | | |
|---|---|---|---|---|
| | 3-α-Maltosylglucose (Koto et al., 1992) | 3-α-Maltosylglucose | 3-α-Maltotriosylglucose | 3-α-Maltotetraosylglucose |
| 1α | 93.2 | 93.2 | 93.2 | 93.2 |
| 1β | 96.9 | 96.9 | 96.9 | 96.9 |
| 2α | 71.1 | 71.1 | 71.1 | 71.1 |
| 2β | 73.8 | 73.8 | 73.8 | 73.8 |
| 3α | 80.5 | 80.5 | 80.4 | 80.4 |
| 3β | 83.1 | 83.1 | 83.1 | 83.1 |
| 4 | 71.0 | 71.0 | 71.0 | 71.0 |
| 5α | 72.2 | 72.2 | 72.2 | 72.2 |
| 5β | 76.6 | 76.6 | 76.6 | 76.6 |
| 6α | 61.2 | 61.2 | 61.1 | 61.1 |
| 6β | 61.6 | 61.5 | 61.5 | 61.5 |
| 1' | 99.8 | 99.8 | 99.8 | 99.8 |
| 2'α | 72.5 | 72.5 | 72.5 | 72.5 |
| 2'β | 72.4 | 72.4 | 72.4 | 72.4 |
| 3' | 74.3 | 74.3 | 74.3 | 74.3 |
| 4'α | 77.7 | 77.7 | 77.8 | 77.7 |
| 4'β | 77.6 | 77.6 | 77.7 | 77.6 |
| 5' | 71.3 | 71.3 | 71.2 | 71.2 |
| 6' | 61.4 | 61.3 | 61.3 | 61.3 |
| 1" | 100.6 | 100.6 | 100.4 | 100.4 |
| 2"α | 72.7 | 72.7 | 72.5 | 72.5 |
| 2"β | | | 72.1 | 72.1 |
| 3" | 73.8 | 73.9 | 74.3 | 74.3 |
| 4" | 70.3 | 70.3 | 77.7 | 77.6 |
| 5" | 73.7 | 73.6 | 71.0 | 71.0 |
| 6" | 61.5 | 61.4 | 61.4 | 61.4 |
| 1''' | | | 100.7 | 100.6 |
| 2'''α | | | 72.7 | 72.5 |
| 2'''β | | | | 72.2 |
| 3''' | | | 73.8 | 74.3 |
| 4''' | | | 70.3 | 77.9 |
| 5''' | | | 73.7 | 71.0 |
| 6''' | | | 61.4 | 61.4 |
| 1'''' | | | | 100.7 |
| 2'''' | | | | 72.7 |
| 3'''' | | | | 73.8 |
| 4'''' | | | | 70.3 |
| 5'''' | | | | 73.7 |
| 6'''' | | | | 61.4 |

**Supplementary Table 4. Effect of metal ions and EDTA on rAoAgtA activity.**

| Chemical | Relative activity (%) |
|---|---|
| Control | 100 ± 6 |
| $Li^{+}$ | 105 ± 3 |
| $K^{+}$ | 102 ± 1 |
| $Mg^{2+}$ | 100 ± 3 |
| $Ca^{2+}$ | 111 ± 3 |
| $Mn^{2+}$ | 110 ± 12 |
| $Fe^{3+}$ | 63 ± 3 |
| $Co^{2+}$ | 92 ± 11 |
| $Cu^{2+}$ | n.d. |
| $Zn^{2+}$ | 56 ± 6 |
| $Al^{3+}$ | 101 ± 4 |
| EDTA | 21 ± 6 |

rAoAgtA activity in the absence of metal ions and EDTA was considered as 100%.

n.d., not detected (no activity).

Data are mean ± standard deviation of three replicates.

#### Supplementary Table 5. rAoAgtA activity in the presence of EDTA and $Ca^{2+}$.

| **Chemicals** | **Relative activity (%)** |
|---|---|
| Control | 100 ± 6 |
| +EDTA | 13 ± 1 |
| +EDTA, $Ca^{2+}$ | 93 ± 5 |

rAoAgtA activity in the absence of EDTA and $Ca^{2+}$ was considered as 100%.

Data are mean ± standard deviation of three replicates.

**Supplementary Table 6. Degradation of various natural glucans by rAoAgtA.**

| Substrate | Glycosidic linkage | Degradation activity |
|---|---|---|
| Corn starch | α-1,4 (primary), α-1,6 | - |
| Potato starch | α-1,4 (primary), α-1,6 | - |
| Soluble starch | α-1,4 (primary), α-1,6 | +, 3.23 ± 0.28 mU/mL |
| Dextran | α-1,4, α-1,6 (primary) | - |
| Pullulan | α-1,4, α-1,6 | - |
| α-1,3-Glucan (bacterial) | α-1,3 | - |
| Nigeran | α-1,3, α-1,4 | - |
| Cellulose | β-1,4 | - |
| Pustulan | β-1,6 | - |
| Laminaran | β-1,3 (primary), β-1,6 | - |

+, degraded; -, not degraded

Data are mean ± standard deviation of three replicates.

#### Supplementary Table 7. Four highly conserved regions of enzymes belonging to the α-amylase family.

| Enzyme | Origin | Sequence in Region I | Sequence in Region II | Sequence in Region III | Sequence in Region IV | NCBI accession number |
|---|---|---|---|---|---|---|
| Agt protein | | | | | | |
| AoAgtA | *Aspergillus oryzae* | 140 DTVINN | 229 GLRIDAAKH | 257 EVLQ | 318 FSENHD | XP_001820542 |
| AnAgtA | *Aspergillus niger* | 142 DTVINN | 231 GLRIDAAKH | 259 EVLQ | 320 FSENHD | XP_003188777 |
| AmyD | *Aspergillus nidulans* | 141 DTVINN | 230 GLRIDAAKH | 258 EVLQ | 319 FSENHD | XP_660912 |
| α-Amylase (Taka-amylase A) | *Aspergillus oryzae* | 138 DVVANH | 223 GLRIDTVKH | 251 EVLD | 313 FVENHD | P0C1B3 |
| CGTase | *Paenibacillus macerans* | 162 DFAPNH | 252 GIRFDAVKH | 285 EWFL | 351 FIDNHD | P31835 |
| Pullulanase | *Klebsiella aerogenes* | 619 DVVYNH | 690 GFRFDLMGY | 723 EGWD | 846 YVSKHD | P07811 |
| Isoamylase | *Pseudomonas amyloderamosa* | 318 DVVYNH | 397 GFRFDLASV | 461 EPWA | 531 FIDVHD | P10342 |
| Branching enzyme | *Escherichia coli* | 335 DWVPGH | 401 ALRVDAVAS | 458 EEST | 521 LPLSHD | P07762 |
| Neopullulanase | *Geobacillus stearothermophilus* | 242 DAVFNH | 324 GWRLDVANE | 357 EIWH | 419 LLGSHD | P38940 |
| Amylopullulanase | *Thermoanaerobacter pseudethanolicus* | 519 DGVFNH | 624 GWRLDVANE | 657 ELWG | 729 LLGSHD | P38939 |
| α-Glucosidase | *Saccharomyces cerevisiae* | 106 DLVINH | 210 GFRIDTAGL | 276 EVAH | 344 YIENHD | P07265 |
| Cyclomaltodextrinase | *Thermoanaerobacter thermohydrosulfuricus* | 238 DAVFNH | 321 GWRLDVANE | 354 EVWH | 416 LIGSHD | A42950 |
| Oligo-1,6-glucosidase | *Bacillus cereus* | 98 DLVVNH | 195 GFRMDVINF | 255 EMPG | 324 YWNNHD | P21332 |
| Dextran glucosidase | *Streptococcus mutans* | 93 DLVVNH | 185 GFRMDVIDM | 231 ETWG | 303 FWNNHD | Q2HWU5 |

The three catalytic residues are shaded in gray. The His residue in Region I conserved among the members of the α-amylase family but replaced with Asn in Agt proteins are highlighted in blue. Numbering of the amino acid sequences of the enzymes starts at N-terminal amino acid of each enzyme including signal peptide.

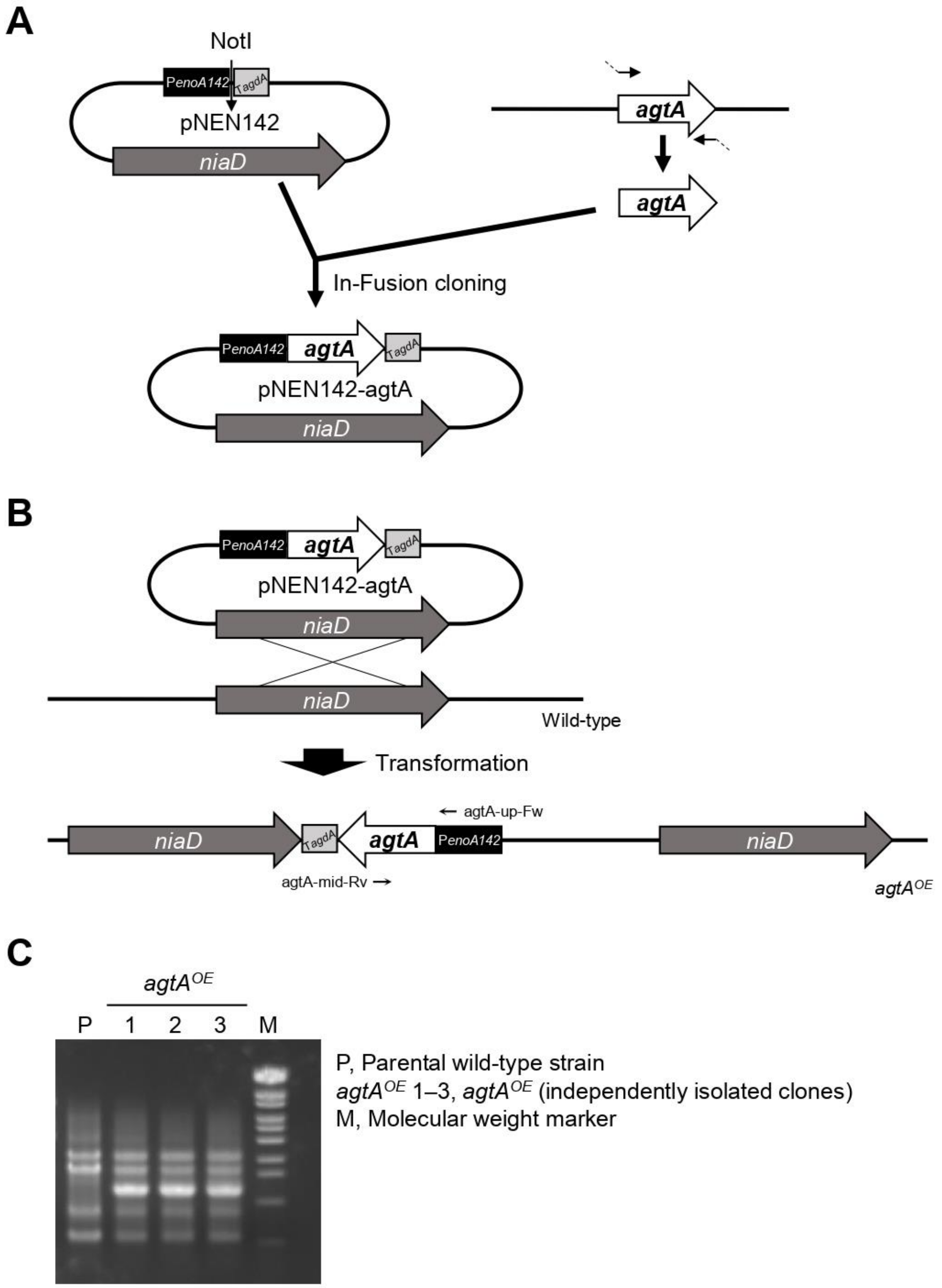


**Supplementary Figure 1. Construction of *agtA*$^{OE}$ strain of *A. oryzae*. (A)** Construction of the pNEN142-agtA plasmid. **(B)** Strategy for *agtA* overexpression. pNEN142-agtA was used to transform the wild-type strain. **(C)** PCR analysis with primers shown in **B** confirmed the integration of the *agtA* overexpression cassette.

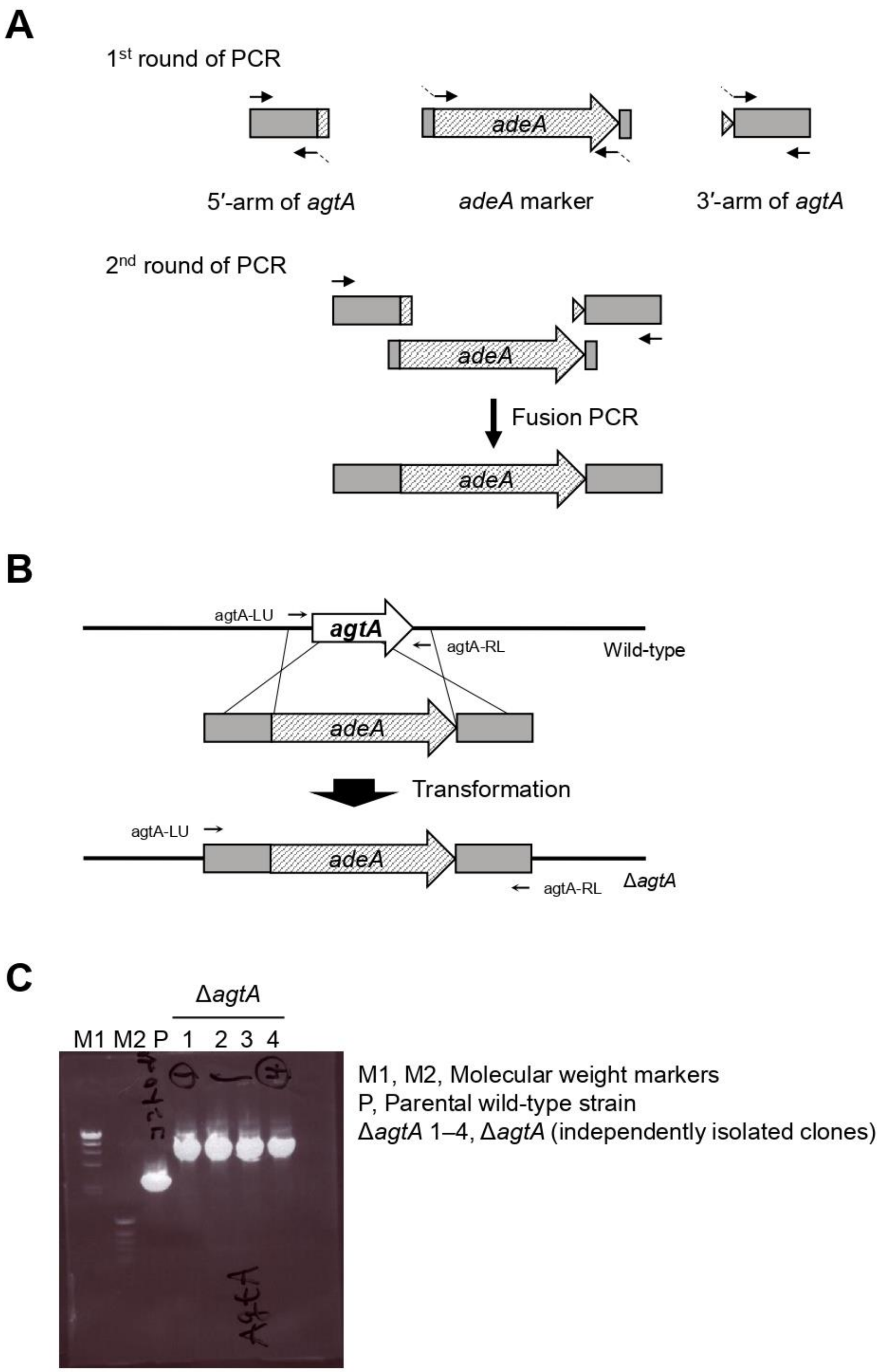


**Supplementary Figure 2. Construction of Δ*agtA* strain of *A. oryzae*. (A)** Construction of the *agtA* disruption cassette. Fragments containing the 5′- and 3′-arms of *agtA* for gene replacement and the *adeA* marker were amplified, and then the three fragments were fused. **(B)** Strategy for *agtA* disruption. The fused cassette was used to transform the wild-type strain. **(C)** PCR analysis of *agtA* gene disruption with primers shown in **B**.

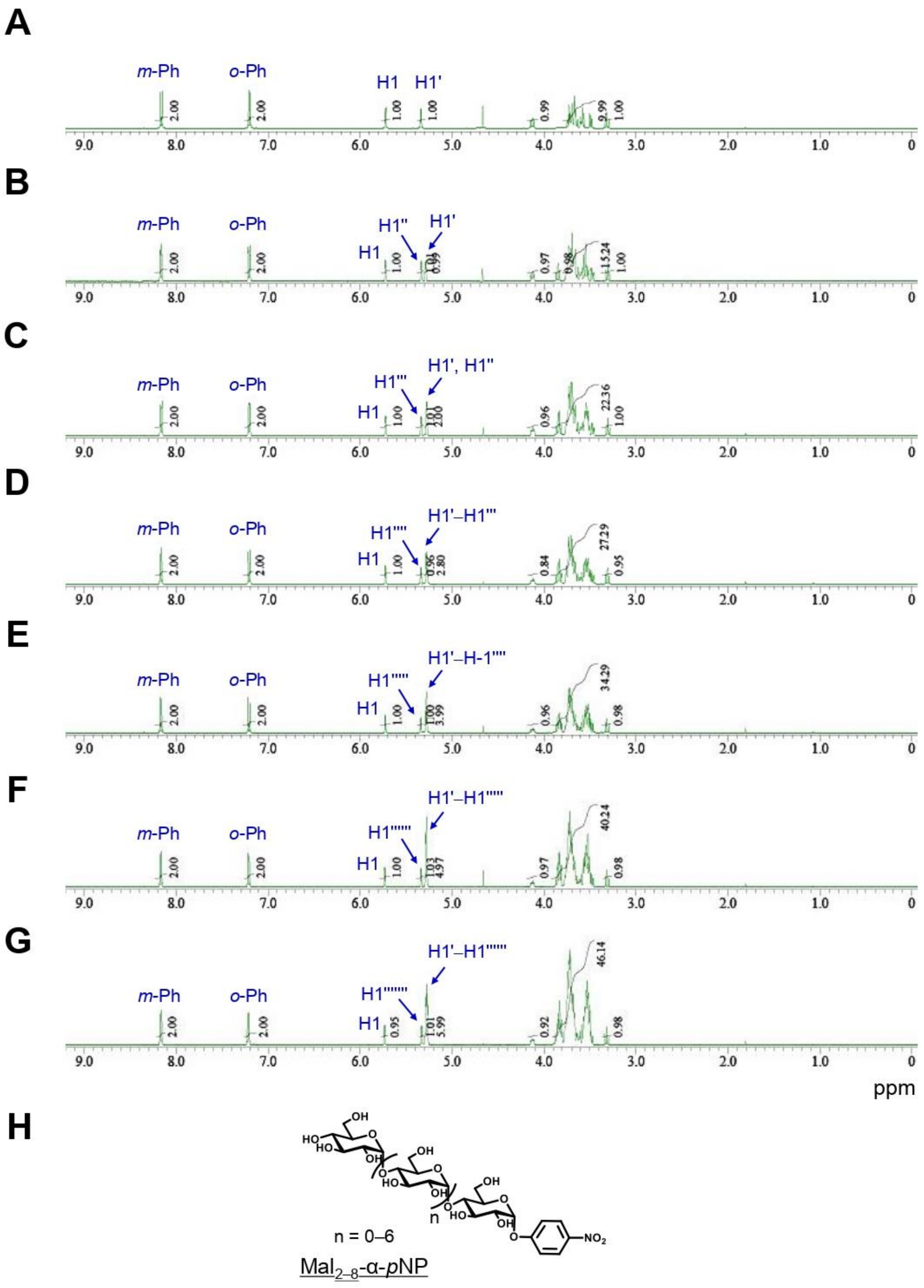


**Supplementary Figure 3. 500 MHz $^1$H-NMR spectra and integral values of Mal$_{2–8}$-α-*p*NP.** Samples: **(A)** Mal$_2$-α-*p*NP; **(B)** Mal$_3$-α-*p*NP; **(C)** Mal$_4$-α-*p*NP; **(D)** Mal$_5$-α-*p*NP; **(E)** Mal$_6$-α-*p*NP; **(F)** Mal$_7$-α-*p*NP; **(G)** Mal$_8$-α-*p*NP. **(H)** Chemical structures of Mal$_{2–8}$-α-*p*NP. Solvent, $D_2O$; temperature, 25°C; concentration, 5 mg/620 μL.

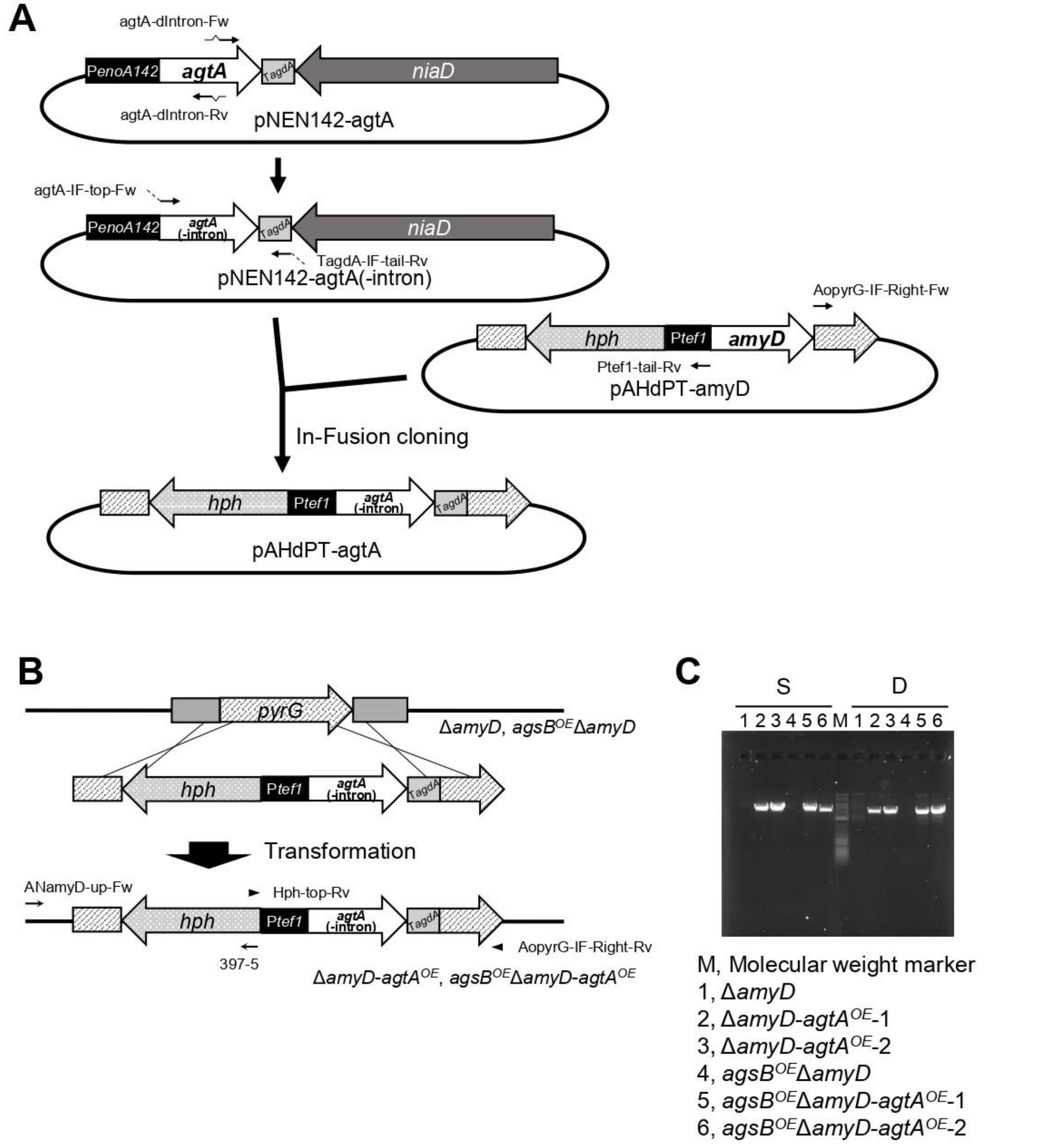


**Supplementary Figure 4. Construction of *agtA*$^{OE}$ strains of *A. nidulans*. (A)** Construction of the pAHdPT-agtA plasmid. The intron of *agtA* was removed from pNEN142-agtA to obtain pNEN142-agtA(-intron). PCR amplification was performed with pNEN142-agtA(-intron) and pAHdPT-amyD as templates, and the two fragments were fused. **(B)** Strategy for overexpression of *agtA* in *A. nidulans*. SacI-digested pAHdPT-agtA was used to transform the Δ*amyD* and *agsB*$^{OE}$Δ*amyD* strains. **(C)** PCR analysis with two pairs of primers shown in **B** (S, arrows; D, arrowheads) confirmed the integration of the *agtA* overexpression cassette.

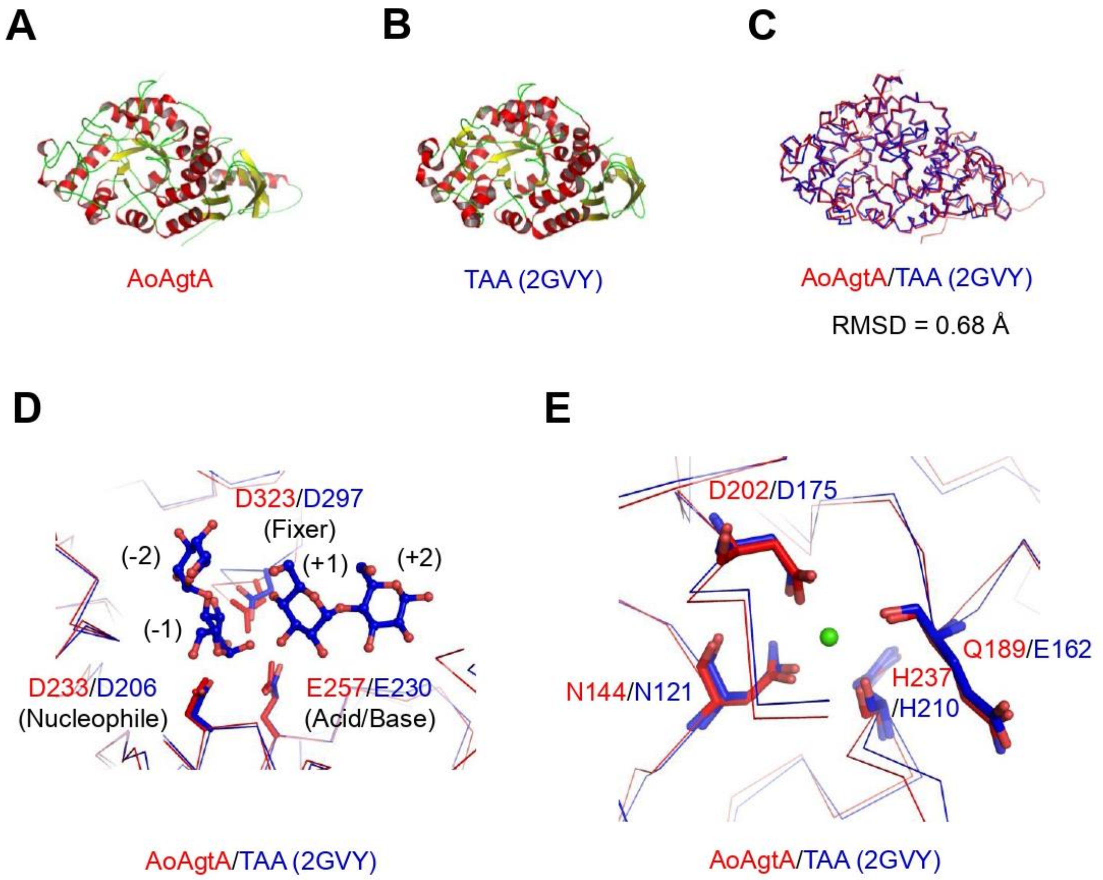


**Supplementary Figure 5. Structure of AoAgtA predicted by AlphaFold2. (A)** The predicted structure of AoAgtA. **(B)** Crystal structure of Taka-amylase A (TAA) (PDB, 2GVY). **(C)** AoAgtA superimposed on TAA. RMSD, root mean square distance between the Cα atoms of 2 aligned residues. **(D)** Catalytic residues and **(E)** $Ca^{2+}$-binding residues of AoAgtA and TAA are conserved. Bound maltose and $Ca^{2+}$ in TAA are shown as ball-and-stick models. Numbers in parentheses indicate the subsites in TAA (**D**). In the panels **D** and **E**, numbering of the amino acid residue starts at the N-terminal of AoAgtA including its signal peptide, and dose that at the N-terminal of TAA without signal peptide.

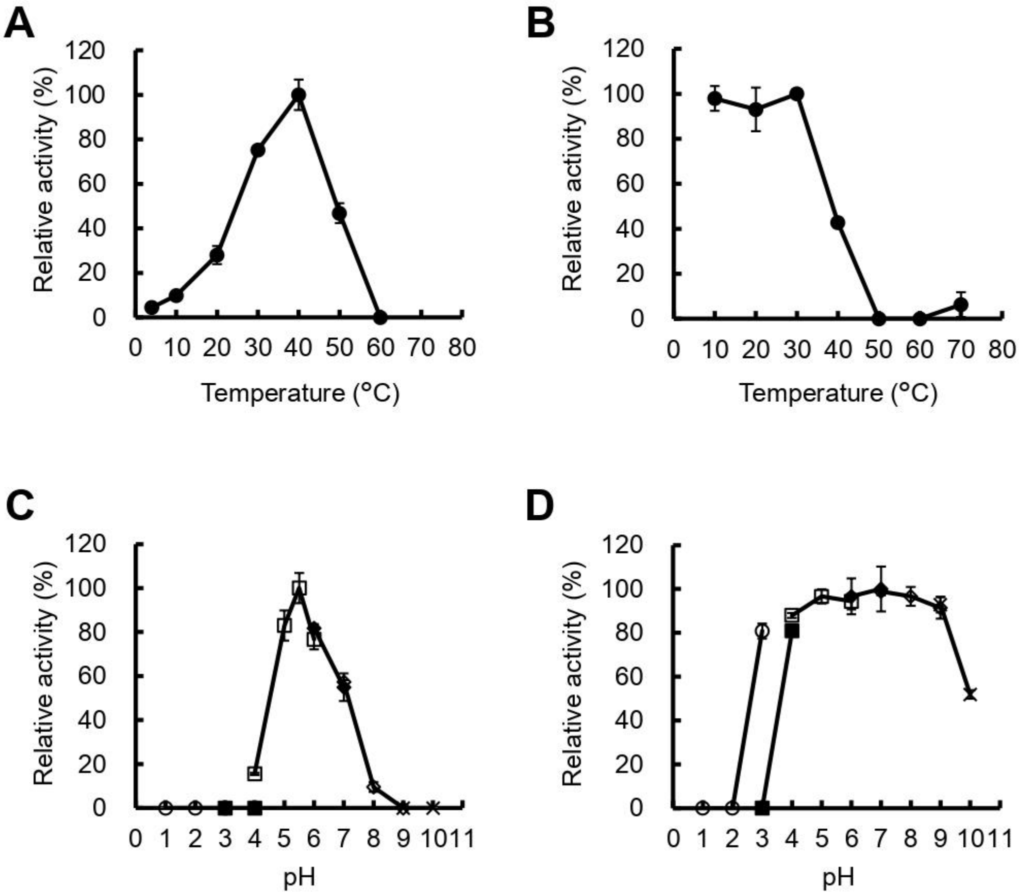


**Supplementary Figure 6. Biochemical characterization of rAoAgtA**. **(A)** Optimal temperature. **(B)** Thermostability. **(C)** Optimal pH. **(D)** pH stability. A mixture (20 μL) containing 1 mM Mal$_5$-α-*p*NP and appropriate rAoAgtA in (**A, B**) 50 mM sodium acetate (Na-Ac) buffer (pH 5.5) or (**C, D**) the indicated buffer and pH was incubated at (**C, D**) 40°C or (**A, B**) the indicated temperature for 10 min. Samples were analyzed by HPLC, and the amount of Mal$_2$-α-*p*NP was quantified to calculate the rAoAgtA activity. Each graph shows relative enzymatic activity, with the highest activity considered 100%. Effect of pH on rAoAgtA activity and stability was examined in different buffers: glycine-HCl (○), citric acid-NaOH (■), Na-Ac (□), MOPS-NaOH (♦), Tris-HCl (◊), and glycine-NaOH (×). Error bars represent the standard deviation of the mean calculated from three replicates.

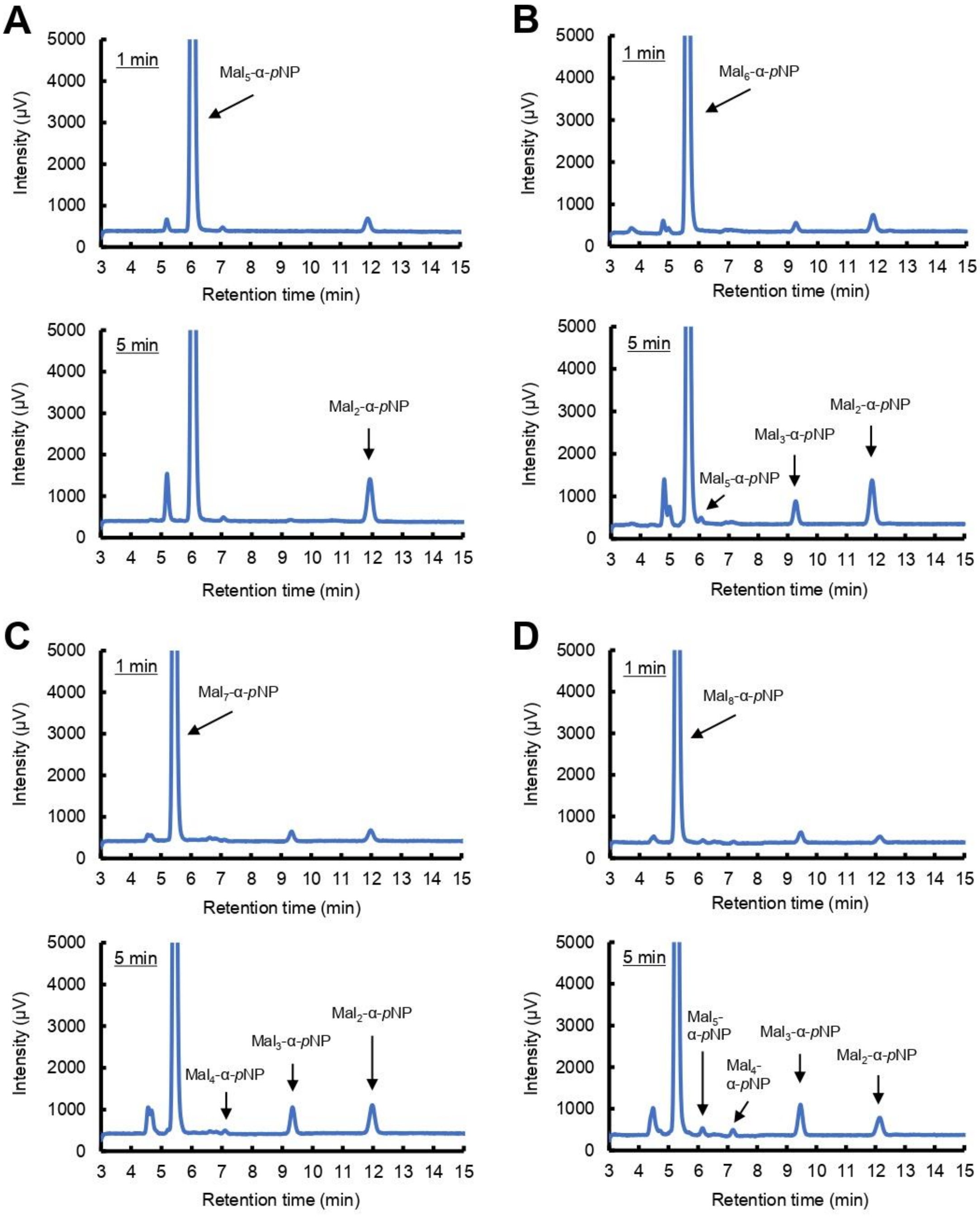


**Supplementary Figure 7. HPLC chromatograms of the substrates and products of Mal$_{5–8}$-α-*p*NP degradation by rAoAgtA.** Substrates: **(A)** Mal$_5$-α-*p*NP; **(B)** Mal$_6$-α-*p*NP; **(C)** Mal$_7$-α-*p*NP; **(D)** Mal$_8$-α-*p*NP. A mixture (20 µL) containing 1.6 mM each substrate and 9.5 mU/mL rAoAgtA in 50 mM Na-Ac buffer (pH 5.5) was incubated at 40°C for 1 or 5 min.

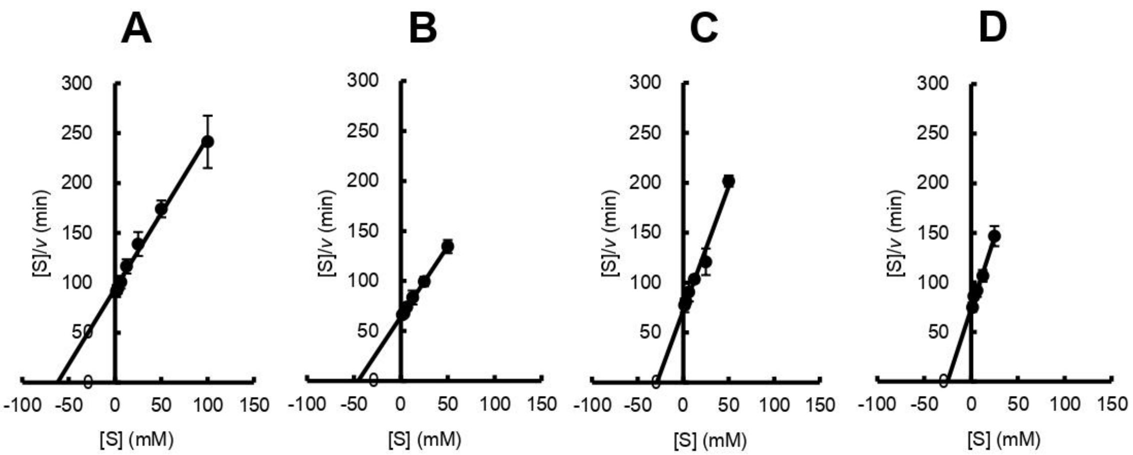


**Supplementary Figure 8. Hanes–Woolf plots for Mal$_{5–8}$-α-*p*NP degradation by rAoAgtA**. Substrates: **(A)** Mal$_5$-α-*p*NP; **(B)** Mal$_6$-α-*p*NP; **(C)** Mal$_7$-α-*p*NP; **(D)** Mal$_8$-α-*p*NP. The kinetic parameters of the reaction were determined by measuring the initial velocity ($v$) (5 min) at each substrate concentration ([S]). A linear transform was used to calculate the Michaelis constant ($K_m$) and maximum velocity ($V_{max}$) of rAoAgtA. Error bars represent the standard deviation of the mean calculated from three replicates.

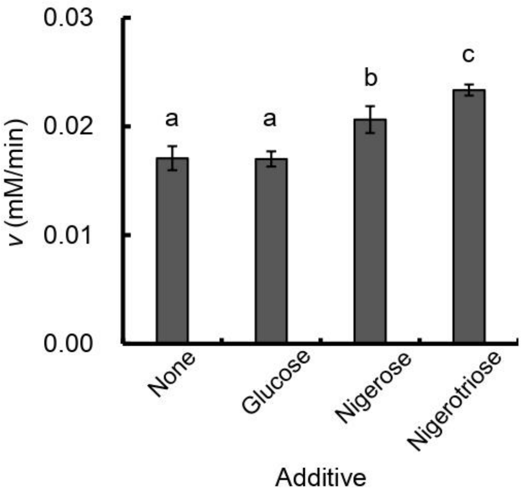


**Supplementary Figure 9**. **The release velocity of $Mal_2$-α-*p*NP from $Mal_5$-α-*p*NP by rAoAgtA with or without nigerooligosaccharides.** A mixture (20 µL) containing 1.6 mM $Mal_5$-α-*p*NP, 9.5 mU/mL rAoAgtA, and 16 mM nigerooligosaccharide or glucose in 50 mM Na-Ac buffer (pH 5.5) was incubated at 40°C for 5 min. Samples were analyzed by HPLC, and the amount of $Mal_2$-α-*p*NP was quantified to calculate its release velocity (*v*). Error bars represent the standard deviation of the mean calculated from three replicates. Different letters above bars indicate significant difference by Tukey's test ($P < 0.01$ none or glucose vs nigerose, none or glucose vs nigerotriose; $P < 0.05$ nigerose vs nigerotriose).

**REFERENCE**

Koto, S., Morishima, N., Shichi, S., Haigoh, H., Hirooka, M., Okamoto, M., et al. (1992). Dehydrative glycosylation using heptabenzyl derivatives of glucobioses and lactose. *Bull. Chem. Soc. Jpn.* 65, 3257–3274. doi: 10.1246/bcsj.65.3257.
